# Turning aging cells into a live vaccine: engineered senescent cancer cells with adjuvant celecoxib for immunotherapy

**DOI:** 10.64898/2025.12.15.694320

**Authors:** Yuewei Wang, Ante Ou, Yanli Luo, Yanrong Gao, Yi Zhang, Linxi Qin, Yongzhuo Huang

## Abstract

The immunoactivation effects of senescent tumor cells are a potential avenue for cancer therapy. They can act as antigen reservoirs for cancer vaccination, but how to maintain strong immunogenicity to induce a robust immunity is underexplored. In this study, we developed an engineered live vaccine composed of hydrogel-encapsulated senescent tumor cells and liposomal celecoxib (STCs+CLX-Lipo@Gel). This vaccine prolongs the in vivo persistence of senescent tumor cells and utilizes liposomal celecoxib (COX2 inhibitor) to promote the recruitment and maturation of dendritic cells (DC). Notably, a single dose can significantly delay melanoma growth by eliciting robust immunity. The vaccine extended the survival of mice with melanoma brain metastases. Moreover, this strategy also demonstrated high efficacy against orthotopic pancreatic tumors. This study presents a comprehensive strategy to boost the immunogenicity of live vaccines by leveraging senescent tumor cells and COX2 inhibition, with treatment efficacy in various tumor models.

**Graphic abstract:** 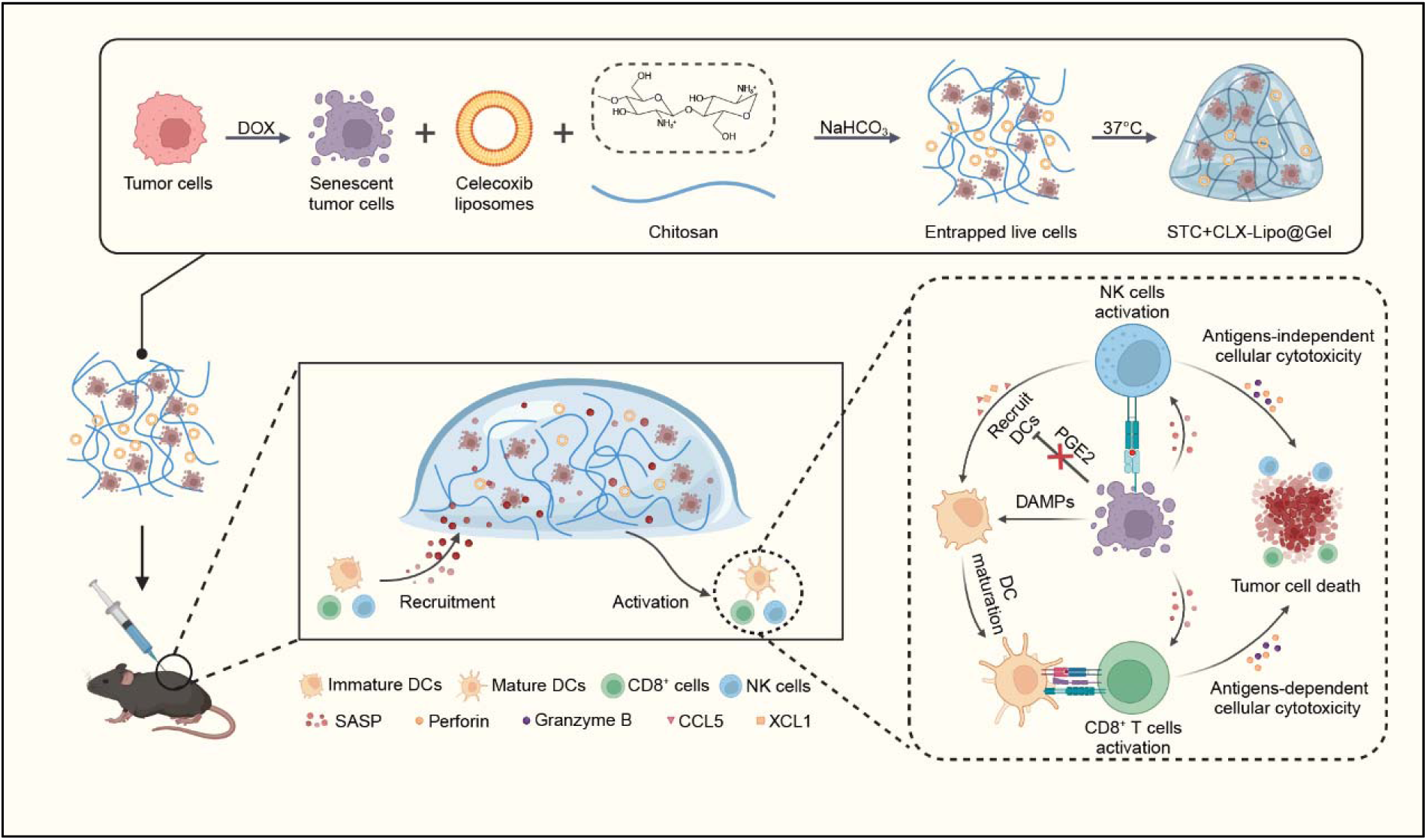

**Graphic summary:** Schematic illustration of the preparation of STC-based live vaccines and the tumor-specific immune responses elicited by the vaccine. Created with BioRender.com (https://BioRender.com/2ind6yh).

## 1 Introduction

Cellular senescence, marked by irreversible cell cycle arrest and a distinctive secretory phenotype, plays a pivotal role in various biological processes, including cancer (Gorgoulis et al., 2019; Marin et al., 2023a). Common anti-cancer therapies—such as radiotherapy, chemotherapy, and immunotherapy—often induce senescence in tumor cells (Gorgoulis et al., 2019; Marin et al., 2023a). While senescent tumor cells (STCs) lose their ability to proliferate, they retain viable state and undergo significant alterations in their surface proteome (Prieto et al., 2023). Importantly, senescent tumor cells persist and continuously produce antigens over time. Besides, STCs exhibit an altered secretome, termed the senescence-associated secretory phenotype (SASP), which includes pro-inflammatory cytokines and chemokines, enabling the recruitment of various immune cells and initiation of multiple immune responses (Lee and Schmitt, 2019; Park et al., 2021). It has been demonstrated that the SASP of senescent tumor cells can recruit and activate CD4^+^ and CD8^+^ T cells, thereby triggering anti-tumor immunity (Iannello et al., 2013; Marin et al., 2023b; Meng et al., 2012; Ruscetti et al., 2020; van Tuyn et al., 2017). Furthermore, senescent tumor cells release damage-associated molecular patterns (DAMPs), which can effectively recruit dendritic cells (DCs) and promote their maturation, acting as immunoadjuvants (Marin et al., 2023a). The multifaceted role of STCs in modulating the immune response highlights their potential as therapeutic targets and as components of immunotherapeutic strategies.

The alterations of proteome and secretome endow STCs with unique properties that can activate both innate and adaptive immune responses, thus being a “rebel force” against tumor growth (Antonangeli et al., 2016; Antonangeli et al., 2019; Chen et al., 2023; Marin et al., 2023a). For example, drug-induced STCs can upregulate ligands for activating receptors on natural killer (NK) cells, such as NKG2D and DNAM-1, thus increasing their susceptibility to recognition and elimination by NK cells (Antonangeli et al., 2016; Antonangeli et al., 2019). Additionally, STCs upregulate MHCI molecules, enhancing cytotoxic T lymphocyte (CD8^+^ T cell) activity (Chen et al., 2023; Marin et al., 2023a). Moreover, the upregulation of IFN-γ receptor IFNGR1 on STCs makes them highly responsive to IFN-γ and type I IFN signaling, and thus STCs typically enhance MHCI antigen processing and presentation, further activating CD8^+^ T cells (Chen et al., 2023; Marin et al., 2023a). Due to their distinctive immunogenic properties and a comprehensive antigen profile, STCs have emerged as promising candidates for live vaccines (Liu et al., 2023; Marin et al., 2023a).

However, there is a primary challenge to overcome. Prostaglandin E2 (PGE2), an essential immunosuppressor secreted by STCs, is prevalent in tumor microenvironment and hampers effective immunoactivation (Bluth et al., 2009; Böttcher et al., 2018; Von Bergwelt-Baildon et al., 2006). PGE2 can inhibit chemokine secretion by NK cells, as well as the proliferation and maturation of DCs, thus diminishing the cytotoxic efficacy of antigen-specific CD8^+^ T cells (Böttcher et al., 2018; Böttcher and Sousa, 2018; Kyrysyuk and Wucherpfennig, 2023; Wculek et al., 2020). To address this issue, we proposed a novel vaccination strategy of utilizing STCs as live vaccines and meanwhile suppressing PGE2 production for augmenting anti-tumor immune responses.

In this work, we developed a STC-based live vaccine and designed a formulation (designated STCs+CLX-Lipo@Gel), in which STCs and liposomal celecoxib were encapsulated into chitosan hydrogel. This design created a niche to prevent STC clearance and extend action duration. Notably, celecoxib is an FDA-approved inhibitor of cyclooxygenase 2 (COX2), and liposomal celecoxib can block COX2/PGE2 pathway to augment the vaccination efficacy. COX2 is a key enzyme controlling PGE2 synthesis, thus serving as a therapeutic target for inhibiting PGE2 production and enhancing immune cell functions (Deng et al., 2022; Jahani et al., 2023). STCs+CLX-Lipo@Gel was expected to harness the antigenicity and adjuvanticity of STCs to stimulate both innate and adaptive immune responses.

## 2 Results

### 2.1 Activation of anti-tumor immunity by senescent tumor cells

Utilizing doxorubicin as a senescence-inducing agent (Marin et al., 2023a), we demonstrated the successful induction of senescence, evidenced by the upregulation of senescence-associated β-galactosidase (SAβG) activity (Figure 1A) and senescence-associated gene expression (Figure S1A), and the increased production of SASPs (e.g., IL-6 and CXCL1) (Figure S1B). STCs exhibited elevated secretion of ATP and HMGB1, and canonical DAMPs, and notably promoted the differentiation of CD11c^+^ MHCII^+^ DCs (Figure 1B and C). It is widely recognized that cellular senescence leads to alterations of the surface proteome and the release of a spectrum of chemokines and pro-inflammatory cytokines, thereby making them susceptible to immune surveillance and immune clearance (Gasek et al., 2021; Hasegawa et al., 2023). Our results revealed a marked upregulated expression of immune cell chemokines (CCL2, CCL5) and antitumor cytokines (IFN-γ, IL-2, IL-12, and IL-15) (Figure S2A and C); these chemokines are crucial for the recruitment and activation of CD8^+^ T cells and NK cells (Böttcher et al., 2018; Marin et al., 2023b; Takasugi et al., 2023). Additionally, the NKG2D ligand proteins (ULBP1, RAE-1) and DNAM-1 ligand protein (CD155), which are pivotal for the activation of NK cells upon receptor binding (Antonangeli et al., 2019), were also significantly elevated (Figure S2B). Furthermore, STCs coculturing with splenic lymphocytes led to a high activation rate of CD8^+^ T cells and NK cells (Figure 1D and E). The secretion of cytotoxic perforin and granzyme B by these immune cells was significantly increased in the STCs-coculturing group (Figure 1F and G), further indicating an enhanced capacity to kill tumor cells. Our results were in agreement with previous studies that highlighted the interaction of STCs with immune cells (Antonangeli et al., 2019; Böttcher et al., 2018; Marin et al., 2023b; Takasugi et al., 2023).

**Figure 1.**
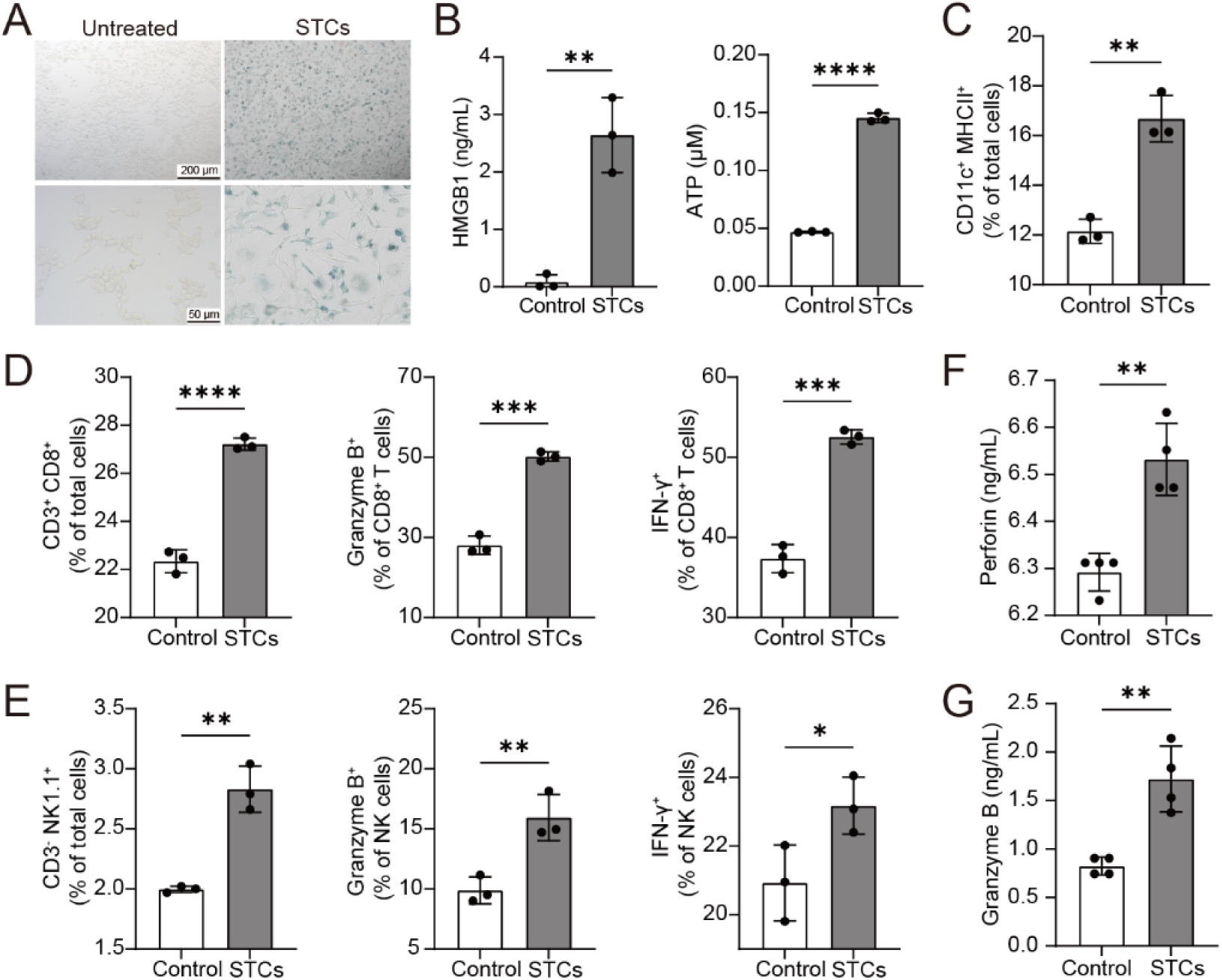
Activated anti-tumor immunity of senescent B16-F10 cells in vitro. (A) Images of senescence-associated β-galactosidase staining of B16-F10 cells (top panel scale bar: 200 μm; bottom panel scale bar: 50 μm). (B) The concentrations of HMGB1 and ATP in the supernatant of untreated cells and STCs. (C)The percentage of CD11c^+^MHCII^+^ DCs after treatment with untreated cells and STCs. (D) The percentage of CD3^+^CD8^+^, CD8^+^Granzyme B^+,^ and CD8^+^IFN-γ^+^T cells after treatment with untreated cells and STCs. (E) The percentage of CD3^-^NK1.1^+^, NK1.1^+^Granzyme B^+,^ and NK1.1^+^IFN-γ^+^ cells after treatment with untreated cells and STCs. The concentrations of (F) Perforin and (G) Granzyme B in the supernatant of untreated cells and STCs (n=4). Statistical significance was assessed by unpaired Student’s t-test. Data were presented as mean ± SD. Statistical significance was determined as *p < 0.05, **p < 0.01, ***p < 0.001, ****p < 0.0001.

In a melanoma model, we further investigated the anticancer efficacy of STCs (Figure 2A). As illustrated in Figure 2B–D and Figure S3, both the died STC group and the STC group significantly inhibited tumor growth, while the STC group exhibited a better efficacy. This reminded us that live STCs could persist releasing antigens and pro-inflammatory cytokines to provide ongoing immune activation signals. By analyzing intratumoral infiltration of immune cell proportions within the primary tumor, the STC group revealed a more prominent anticancer effect, evidenced by a significant increase of CD4^+^ and CD8^+^ T cells (Figure 2E). Moreover, the ratio of CD8^+^ Granzyme B^+^ and CD8^+^ Ki67^+^ T cell subsets was significantly upregulated in the STC group, significantly higher than that in the died STC group (Figure S4A). A significantly increased proportion of effector memory T cells (TEMs), NK cells, and granzyme B^+^ NK cells in the spleens were also observed in the STC group (Figure 2F and G). These experimental results revealed the efficient activation of the anticancer T cell immunity by both the died STC group and the STC group, while the STC group exhibited better efficacy, possibly due to the prolonged action duration of the live STCs compared to the died STCs.

**Figure 2.**
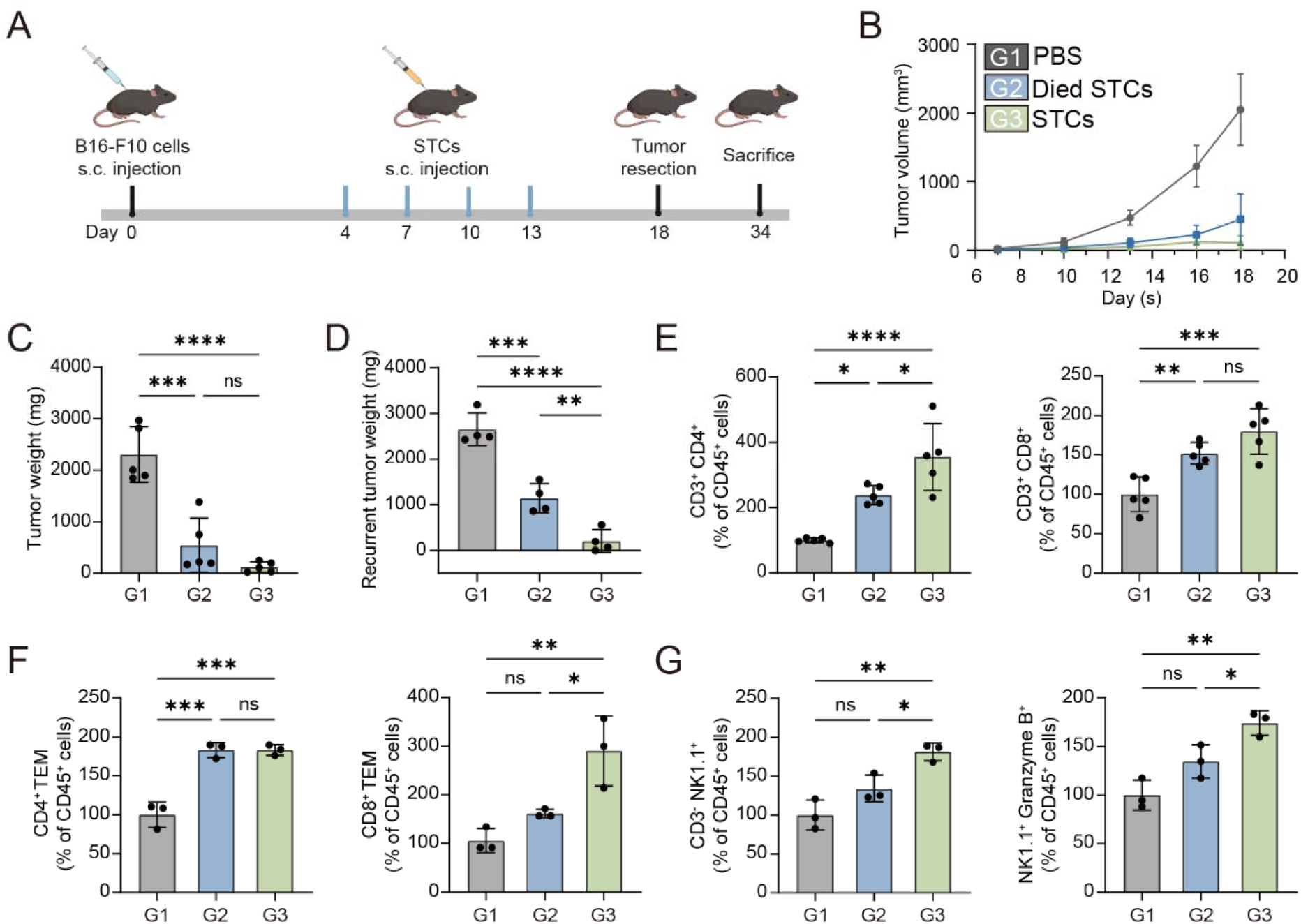
Immunization with senescent B16-F10 cells promoted anticancer immunity. (A) Schedule of STCs treatment in a recurrent melanoma model. Created with BioRender.com (https://BioRender.com/ttybc4v). (B) The average tumor volume change of mice treated with different groups (n=5). (C) The tumor weight of the primary tumors (n=5). (D) The tumor weight of recurrent tumors (n=4). (E) The percentage of CD3^+^CD4^+^ and CD3^+^CD8^+^ T cells within tumors treated with different groups (n=5). (F) The percentage of CD4^+^ and CD8^+^ T cells within spleens of mice treated with different groups (n=3). (G) The percentage of CD3^-^NK1.1^+^ and NK1.1^+^Granzyme B^+^ cells in the spleens of mice treated with different groups (n=3). Statistical significance was assessed by one-way ANOVA and Tukey multiple comparisons tests. Data were presented as mean ± SD. Statistical significance was determined as *p < 0.05, **p < 0.01, ***p < 0.001, ****p < 0.0001.

### 2.2 Promotion of DCs recruitment by celecoxib

Of note, intratumoral infiltration of DCs following STC treatment did not significantly increase compared to the died STC group (Figure S4B). This could be accounted for that DC recruitment into tumors is typically regulated by NK cells by secreting chemokines such as CCL5 and XCL1 (Böttcher et al., 2018). However, this process can be disrupted by STC-produced PGE2 which inhibits NK cells’ activity and production of chemokines (Böttcher et al., 2018; Böttcher and Sousa, 2018). Upregulated COX2 in cellular senescence typically leads to increased PGE2 production (Gorgoulis et al., 2019). Therefore, we proposed the application of liposomal celecoxib (COX2 inhibitor) could reduce PGE2 production and promote DC recruitment.

As shown in Figure 3A, S12, the STCs indeed exhibited higher COX2 expression levels compared to the control cells without senescence. With celecoxib treatment, STCs demonstrated a remarkable decrease in COX2 expression and PGE2 production (Figure 3B–C), but without an impact on the anticancer SASP factors (e.g., IL-6 and TNF-α) (Figure S5).

**Figure 3.**
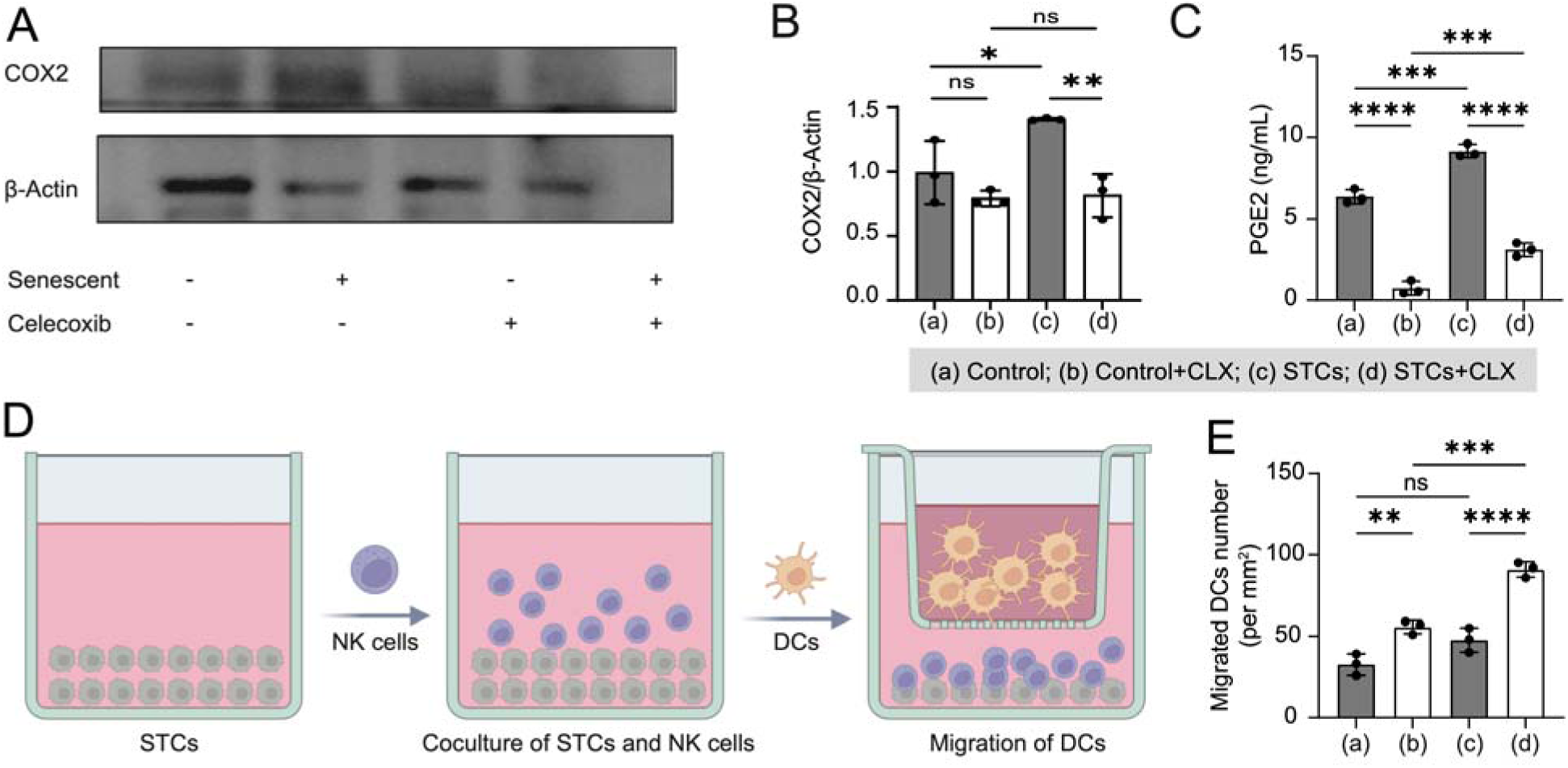
Celecoxib promoted the DCs recruitment. (A–B) COX2 expression in senescent and normal B16-F10 cells with or without celecoxib treatment detected by western blotting and statistical analysis. (C) PGE2 concentrations in the supernatant of different treatments. (D) Schematic diagram of Transwell coculture system. Created with BioRender.com (https://BioRender.com/ttybc4v). (E) The quantitative analysis of migrated DCs after different treatments. Statistical significance was assessed by one-way ANOVA and Tukey multiple comparisons tests. Data were presented as mean ± SD (n=3). Statistical significance was determined as ns (not significant, p > 0.05), **p < 0.01, ***p < 0.001, ****p < 0.0001.

The ability to recruit DCs was evaluated through Transwell experiments after coculturing NK cells with different pretreatment (Figure 3D). STCs activated NK cells more effectively than the Control group, but the amount of migrating DCs recruited by the NK cells was not significantly increased (Figure 3E). By contrast, the NK cells activated by the celecoxib-treated STCs markedly increased the recruitment of DCs (Figure 3E). Similarly, such an effect was also achieved by using another COX2 inhibitor meloxicam, and addition of exogenous PGE2 reversed the enhanced DC-recruitment effect of both inhibitors (Figure S6). These results collectively demonstrated that coapplication of celecoxib with STCs enhanced the NK cell activation and DC recruitment efficiency.

### 2.3 Characterization of STCs+CLX-Lipo@Gel

The preparation of chitosan hydrogel incorporating cells and liposomes was delineated in Figure 4A. Due to the limited solubility of celecoxib in aqueous solutions, we formulated celecoxib liposomes (referred to as CLX-Lipo) for enhanced biocompatibility to the hydrogel matrix. The CLX-Lipo exhibited a spherical morphology with a size of approximately 60 nm and negative zeta potential (Figure 4B–D); the drug encapsulation efficiency was 73.5% and the drug-loading capacity was 2% (Table S1).

**Figure 4.**
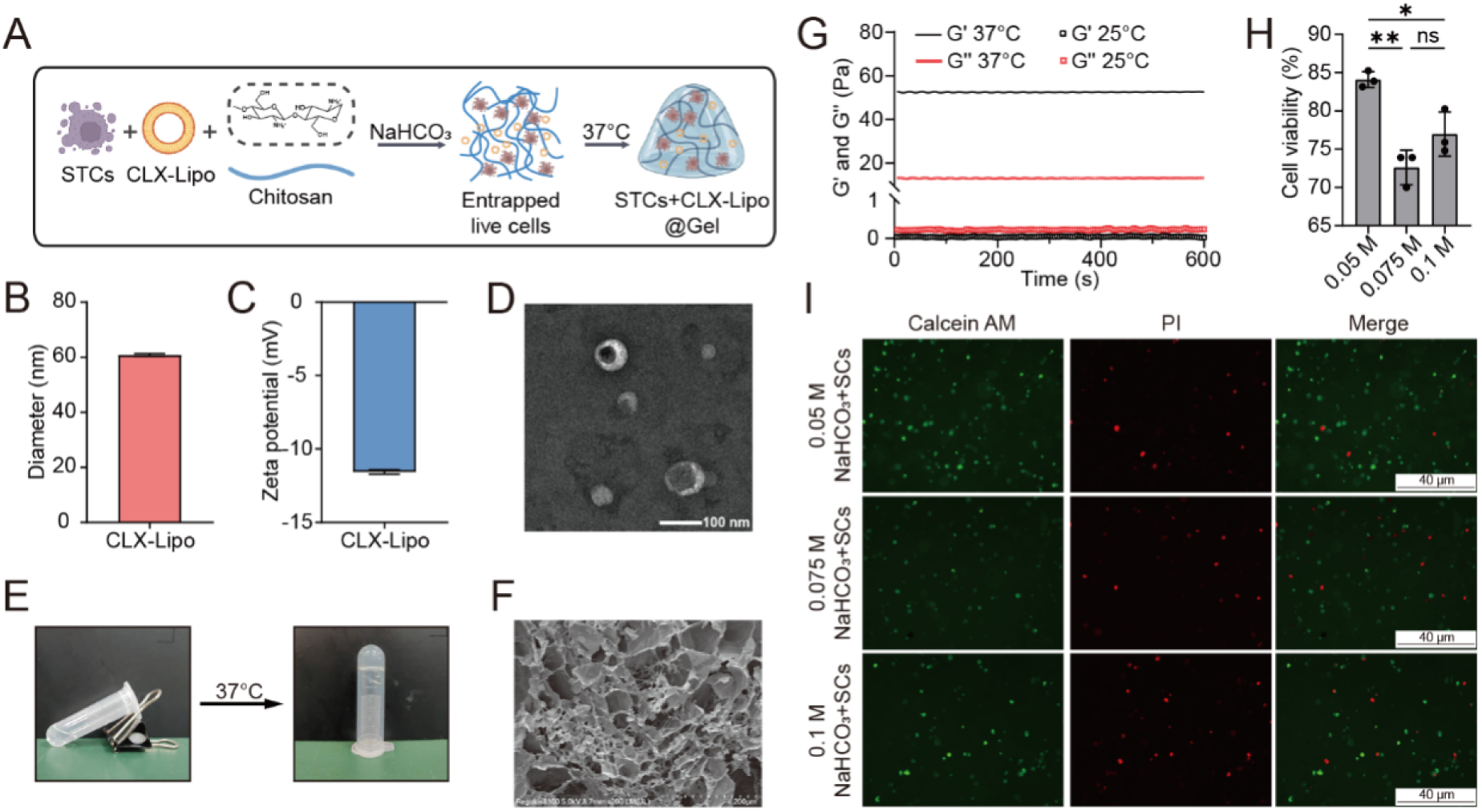
Preparation and characterization of STCs+CLX-Lipo@Gel. (A) Schematic diagram of the preparation process for STCs+CLX-Lipo@Gel. Created with BioRender.com (https://BioRender.com/nx2mmfx). (B) Particle size and (C) zeta potential of CLX-Lipo. (D) Transmission Electron Microscopy image of CLX-Lipo (scale bar: 100 nm). (E) Photograph of STCs+CLX-Lipo@Gel post-gelatinization. (F) Scanning Electron Microscopy (SEM) image of STCs+CLX-Lipo@Gel. (G) Rheological evaluation of STCs+CLX-Lipo@Gel over time. (H) Quantitative analysis of cell viability for STCs encapsulated within chitosan hydrogel and (I) representative fluorescence images for Live/Dead staining (green: Calcein-AM; red: PI; scale bar: 40 μm). Statistical significance was assessed by one-way ANOVA and Tukey multiple comparisons tests. Data were presented as mean ± SD (n=3). Statistical significance was determined as ns (not significant, p > 0.05), *p < 0.05, **p < 0.01.

An injectable, temperature-sensitive chitosan gel was prepared using phosphate buffer (PB) containing sodium bicarbonate (NaHCO_3_) as the gelling agent. NaHCO_3_ neutralizes the protons of amino groups in chitosan chains, thereby reducing interchain repulsions (Ahmadi and de Bruijn, 2008; Assaad et al., 2015; Monette et al., 2016). By reaching a balance of electrostatic attractions and interchain repulsions, gelation was formed by the entangled chains. An increase in temperature accelerates proton transfer, endowing this hydrogel with temperature-sensitive properties (Ahmadi and de Bruijn, 2008; Assaad et al., 2015; Monette et al., 2016). The gelation times at a temperature of 37°C with varying

NaHCO_3_ concentrations (0.05 M, 0.075 M, and 0.1 M) were all in 1 h (Figure 4E and Table S2). The thus-formed CS@Gel exhibited a reticulated porous structure, with pore size ranging from 5 to 100 μm (Figure 4F). Rheological analysis revealed that the storage modulus (G’) of the CS@Gel was significantly higher than the loss modulus (G’’), indicating a robust elastic property (Figure 4G). Cell viability remained at 85% after 14 days of culture with the CS@Gel (Figure 4H and I). Given that the 0.05 M group demonstrated the highest cell viability, we employed a NaHCO_3_ concentration of 0.05 M to establish the hydrogel system in subsequent experiments.

### 2.4 DCs and T cells activation by STCs+CLX-Lipo@Gel

We further investigated the synergistic effect of the hydrogel system in activating DCs (Figure 5A). The flow cytometry result showed that the percentage of MHCII^+^ DCs was 82% in the STCs+CLX-Lipo@Gel-treated group, significantly higher than that in the STCs+CLX-Lipo and STCs-treated groups (Figure 5B and C). Conventional type 1 dendritic cells (cDC1) are efficient at presenting exogenous antigens, such as tumor antigens, to CD8^+^ T cells (Böttcher et al., 2018; Böttcher and Sousa, 2018; Ferris et al., 2020). They are pivotal in eliciting cytotoxic effector T-cell responses and orchestrating anti-cancer immune responses (Böttcher and Sousa, 2018). As illustrated in Figure 5D and E, the proportion of cDC1 cells in the PBS group was low (0.3%). After stimulation with STCs, CLX-Lipo-treated STCs, or STC+CLX-Lipo@Gel, the proportion of cDC1 cells in mature DCs was 2.3%, 7.8%, and 14.1%, respectively. These results indicated that STCs+CLX-Lipo@Gel exhibited an improved effect on cDC1 activation, serving as a potent immunoadjuvant.

**Figure 5.**
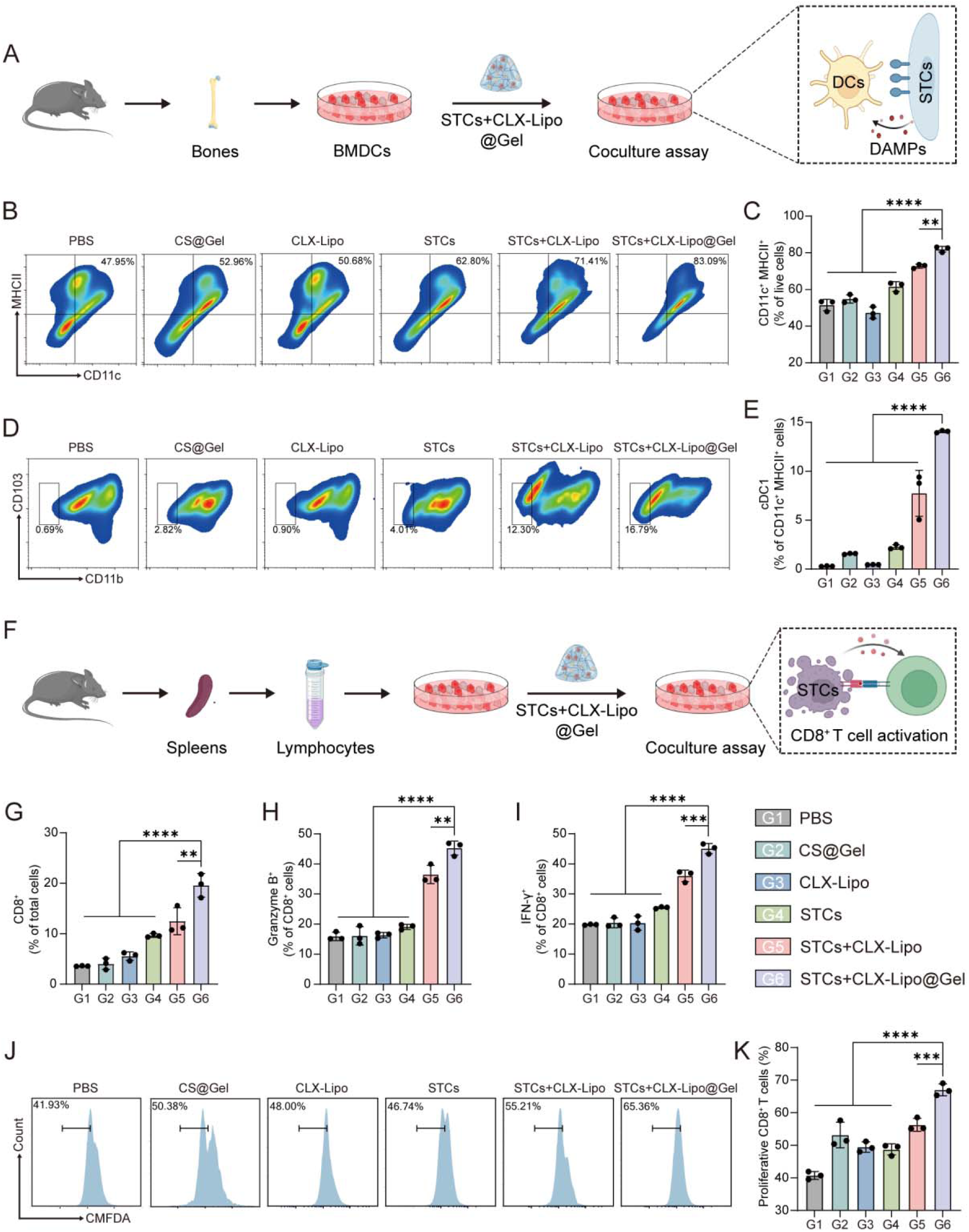
STCs+CLX-Lipo@Gel promoted maturation of DCs and activation of CD8^+^ T cells. (A) Schematic diagram of coculture of STCs+CLX-Lipo@Gel and BMDCs. (B) Flow cytometry analysis and (C) quantitative analysis of the percentage of CD11c^+^ MHCII^+^ cells after various treatments. (D) Flow cytometry analysis and (E) quantitative analysis of the percentage of cDC1 in matured DCs after various treatments. (F) Schematic diagram of coculture of STCs+CLX-Lipo@Gel and splenic lymphocytes. The percentage of (G) CD8^+^ T cells, (H) Granzyme B^+,^ and (I) IFN-γ^+^ in CD8^+^ T cells after various treatments. (J) Flow cytometry histograms depicting CMFDA fluorescence intensity of CD8^+^ cells. (K) Quantitative analysis of the percentage of proliferative CD8^+^ T cells after various treatments. Statistical significance was assessed by one-way ANOVA and Tukey multiple comparisons tests. Data were presented as mean ± SD (n=3). Statistical significance was determined as, **p < 0.01, ***p < 0.001, ****p < 0.0001.

In a coculture model involving T cells and STCs+CLX-Lipo@Gel, there was a very high count of CD8^+^ T cells, approximately 5.4 times than that of the PBS control group (Figure 5F and G), and there was also a high proportion of Granzyme B^+^ and IFN-γ^+^CD8^+^ T cells, reaching approximately 45% (Figure 5H and I). In addition, the STCs+CLX-Lipo@Gel treatment also increased the proportion of the proliferative CD8^+^ T cells, consistent with the findings above (Figure 5J and K).

### 2.5 Restoration of the functionality of the NK-DC axis by STCs+CLX-Lipo@Gel

NKG2D and NKp46, the essential surface receptors on NK cells, can recognize and bind to infected or malignant cells, thereby activating NK cells and facilitating the elimination of target cells (Antonangeli et al., 2016; Kyrysyuk and Wucherpfennig, 2023). We assessed the activation of NK cells through a coculture model (Figure 6A). Compared to the PBS group, both the STC group and the STCs+CLX-Lipo group showed a significant increase in the proportion of NKG2D^+^ and NKp46^+^ NK cells, while the STCs+CLX-Lipo@Gel group exhibited the highest efficacy (Figure 6B and C; Figure S7A and B). Besides, coculturing with STCs+CLX-Lipo@Gel resulted in a substantial increase in the proportion of proliferating NK cells (Figure 6D; Figure S7C). Consequently, the STCs+CLX-Lipo@Gel treatment markedly enhanced the cytotoxic capacity of NK cells against the tumor cells, as illustrated in Figure 6E.

**Figure 6.**
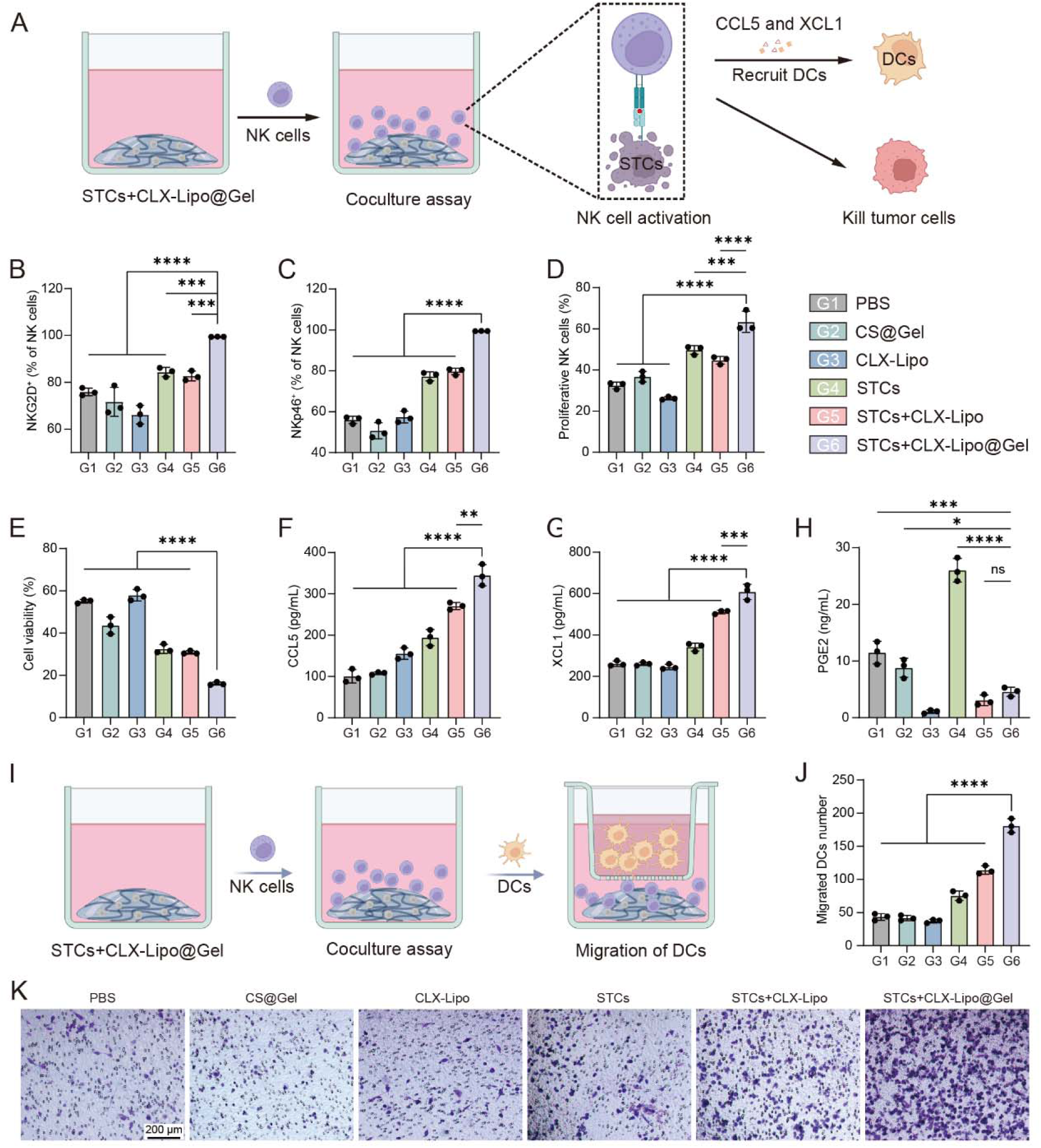
STCs+CLX-Lipo@Gel activated NK cells and restored the functionality of the NK-DC axis. (A) Schematic diagram of coculture of STCs+CLX-Lipo@Gel and NK cells. The (B) NKG2D^+^ and (C) NKp46^+^ cell percentages in NK cells following various treatments. (D) Quantitative analysis of the percentage of proliferative NK cells after various treatments. (E) The ability of NK cell-mediated killing B16-F10 cells after various treatments detected by CCK8 assay. The concentrations of (F) CCL5, (G) XCL1, and (H) PGE2 in the supernatant of NK cells after various treatments. (I) Schematic diagram of Transwell coculture system. Created with BioRender.com (https://BioRender.com/ttybc4v). (J) The quantitative analysis and (K) representative images of migrated DCs after different treatments (scale bar: 200 μm). Statistical significance was assessed by one-way ANOVA and Tukey multiple comparisons tests. Data were presented as mean ± SD (n=3). Statistical significance was determined as ns (not significant, p > 0.05), *p < 0.05, **p < 0.01, ***p < 0.001, ****p < 0.0001.

Notably, inhibition of PGE2 production can reinstall the NK-DC axis and thus promote the recruitment of DCs (Böttcher et al., 2018; Böttcher and Sousa, 2018). Our study revealed that coculture with STCs+CLX-Lipo@Gel led to a marked increase in CCL5 and XCL1 secretion by NK cells (Figure 6F and G), as a result of the significantly reduced levels of PGE2 (Figure 6H). The Transwell assay indicated that the STC group did not promote DC migration, while the STCs+CLX-Lipo@Gel group significantly enhanced DC recruitment, consistent with the aforementioned findings on NK cell chemokine secretion (Figure 6I–K).

### 2.6 In vivo retention of STCs+CLX-Lipo@Gel

DiR-labeled STCs were utilized to prepare STCs@Gel that were subcutaneously injected into mice to investigate the in vivo retention. As shown in Figure 7A and B, the fluorescence intensity of the STC group gradually diminished after subcutaneous injection, reaching only 1/5 of the initial intensity by day 7. By contrast, the STCs@Gel group displayed no substantial decline in fluorescence intensity by the 7th day, with the average fluorescence intensity on day 14 remaining substantial (Figure 7B). These results indicated that STCs encapsulated within a gel could prolong the in vivo retention of STCs.

**Figure 7.**
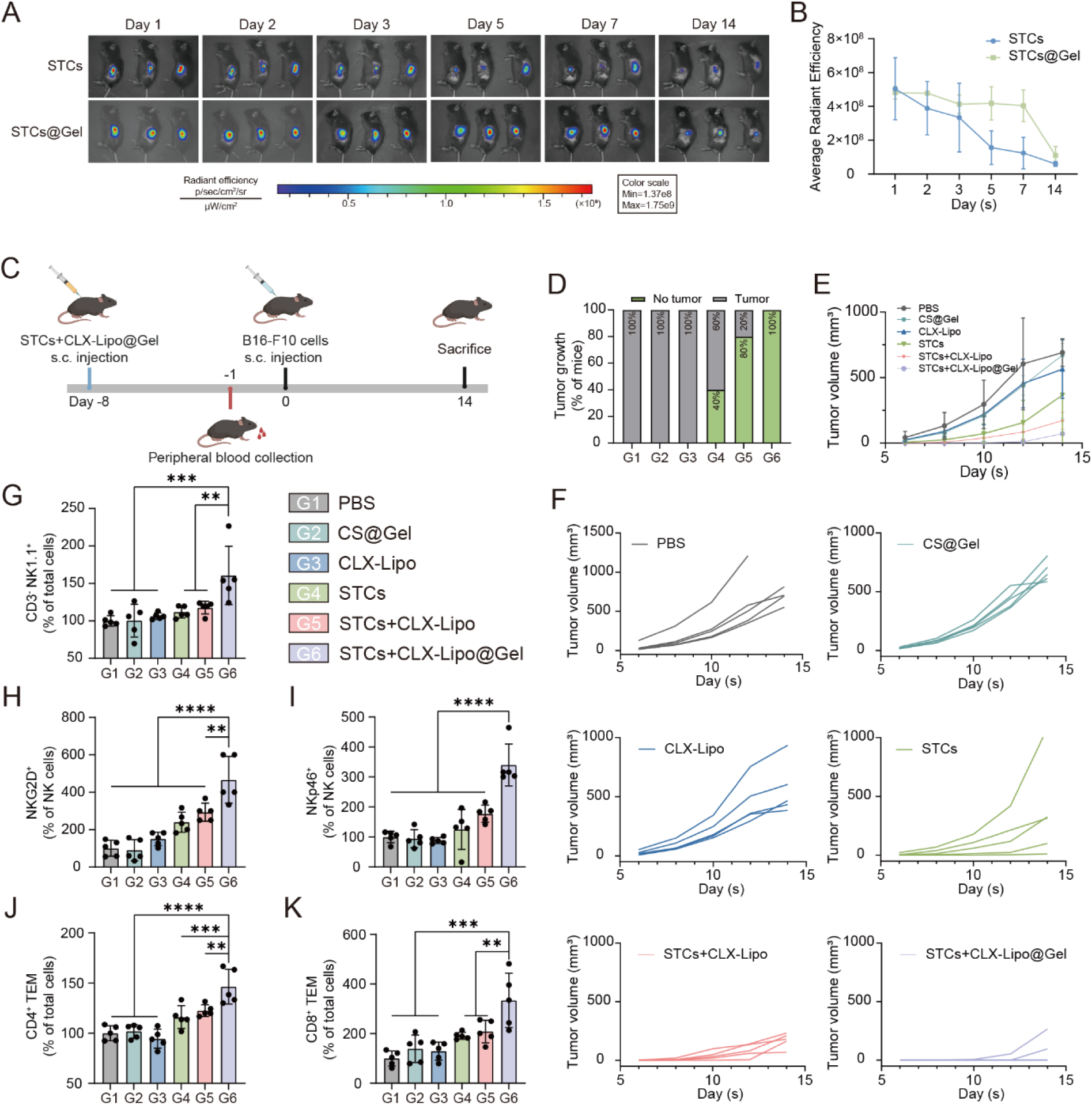
In vivo biodistribution, prophylactic and immunoregulatory effects of STCs+CLX-Lipo@Gel. (A) Representative IVIS images at different time points of DiR-labeled STCs encapsulated in chitosan hydrogel and (B) semi-quantitative analysis of fluorescent signal (n=3). (C) Schedule of various treatments in a melanoma challenge model. Created with BioRender.com (https://BioRender.com/ttybc4v). (D) Tumor-free rate of different groups on day 7. (E) The average tumor growth curve of different groups. (F) The individual tumor growth of each group. (G) The percentage of CD3^-^ NK1.1^+^ cells in peripheral blood after various treatments. The percentage of (H) NKG2D^+^ and (I) NKp46^+^ cells in NK cells in peripheral blood after various treatments. The percentage of (J) CD4^+^ and (K) CD8^+^ effector memory T cells (TEMs) in spleens after various treatments. Statistical significance was assessed by one-way ANOVA and Tukey multiple comparisons tests. Data were presented as mean ± SD (n=5). Statistical significance was determined as *p < 0.05, **p < 0.01, ***p < 0.001, ****p < 0.0001.

### 2.7 In vivo prophylactic efficacy of STCs+CLX-Lipo@Gel

A tumor challenge study was conducted in C57BL/6 mice to evaluate the prophylactic efficacy of the pre-administered STCs+CLX-Lipo@Gel against melanoma development (Figure 7C). By the 7th day, the STCs+CLX-Lipo@Gel group exhibited no observable tumors in any of the mice (Figure 7D), and showed an average tumor volume of only 181 mm^3^ on the 14th day, and three mice did not develop tumors (Figure 7E–F). By contrast, the tumor growth was rapid in other groups, with an average tumor volume exceeding 500 mm^3^ by the 14th day (Figure 7E–F).

To evaluate the immune-stimulatory effect of STCs+CLX-Lipo@Gel, we collected the blood and spleen samples for subsequent analysis. Compared to the PBS group, the STCs+CLX-Lipo@Gel group demonstrated a significant increase in both the amount of NK cells and the proportion of activated NK cells (NKG2D^+^ and NKp46^+^) in the blood of mice (Figure 7G–I). Additionally, there was also a significant increase in the proportions of both CD4^+^ TEMs and CD8^+^ TEMs in the spleens of mice receiving STCs+CLX-Lipo@Gel (Figure 7J–K).

### 2.8 Antitumor effect of STCs+CLX-Lipo@Gel in the subcutaneous melanoma model

The therapeutic efficacy of STCs+CLX-Lipo@Gel was further investigated in a melanoma mouse model (Figure 8A). A single treatment was administered to mice on the 11th day (Figure 8A). As illustrated in Figure 8B and C, the STCs+CLX-Lipo@Gel group exhibited the smallest tumor volume, revealing a significant tumor suppression efficacy (inhibition rate of 94.3%) (Figure 8D and Figure S8).

**Figure 8.**
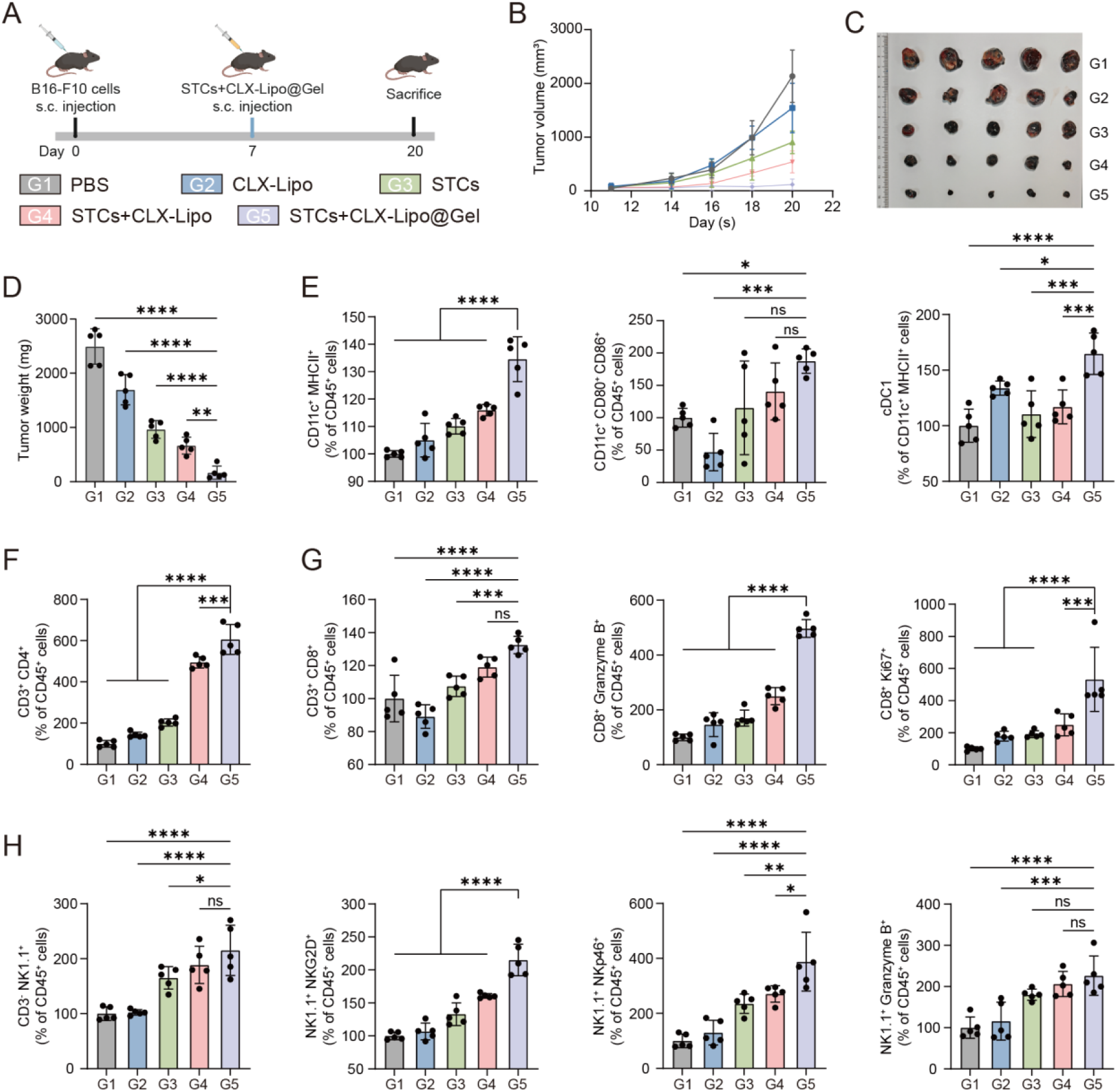
Antitumor and immune-stimulating effects of STCs+CLX-Lipo@Gel in the subcutaneous melanoma model. (A) Schedule of various treatments in subcutaneous melanoma model. Created with BioRender.com (https://BioRender.com/ttybc4v). (B) Tumor growth curve, (C) tumor image, and (D) tumor weight of different groups. (E) The percentage of matured DCs (CD11c^+^ MHCII^+^ and CD11c^+^ CD80^+^ CD86^+^) and cDC1 in peritumoral lymph nodes after various treatments. (F) The percentage of CD4^+^ T cells in tumor tissues after various treatments. (G) The percentage of CD8^+^ T cells and activated CD8^+^ T cells (CD8^+^ Granzyme B^+^) proliferative CD8^+^ T cells (CD8^+^ Ki67^+^) in tumor tissues after various treatments. (H) The percentage of NK cells (CD3^-^ NK1.1^+^), and activated NK cells (NK1.1^+^ NKG2D^+^, NK1.1^+^ NKp46^+,^ and NK1.1^+^ Granzyme B^+^) in tumor tissues after various treatments. Statistical significance was assessed by one-way ANOVA and Tukey multiple comparisons tests. Data were presented as mean ± SD (n=5). Statistical significance was determined as *p < 0.05, **p < 0.01, ***p < 0.001, ****p < 0.0001.

At the experimental endpoint, the lymph nodes and tumor tissues were harvested for flow cytometric analysis of immune cells. The STCs+CLX-Lipo@Gel treatment significantly increased the proportion of mature DCs in the lymph nodes and promoted cDC1 recruitment to the lymph nodes adjacent to the tumors (Figure 8E). The STCs+CLX-Lipo@Gel group also showed a significant increase in the proportion of CD4^+^ and CD8^+^ T cells in the tumor tissue compared to the PBS group (Figure 8F and G). Specifically, there was a notable increase in the proportion of the CD8^+^ Ki67^+^ cell subset, suggesting the enhanced proliferation of intratumoral CD8^+^ T cells (Figure 8G). Concurrently, the proportion of CTLs (CD8^+^ Granzyme B^+^) increased significantly in the STCs+CLX-Lipo@Gel group, approximately 5 times higher than that of the PBS group and 2 times higher than that of the STCs+CLX-Lipo group (Figure 8G). Analysis of intratumoral NK cells showed a significant increase in their proportion following treatment with STCs+CLX-Lipo@Gel (Figure 8H). The amount of activated NK cells (NKG2D^+^ and NKp46^+^) also increased significantly, about 2 and 4 times higher than that of the PBS group, respectively (Figure 8H). Accordingly, the proportion of cytotoxic effector NK cells (Granzyme B^+^) also increased significantly (Figure 8H). These results indicated that STCs+CLX-Lipo@Gel treatment could significantly activate anti-tumor immunity.

### 2.9 Antitumor effect of STCs+CLX-Lipo@Gel in various tumor models

Melanoma brain metastases have emerged as a prevalent and formidable clinical challenge, often culminating in the demise of patients as a direct consequence of the disease (Oliva et al., 2018). We undertook an investigation to ascertain whether this vaccine could mitigate the development of melanoma brain metastases. The mice were vaccinated as depicted in Figure 9A, and the efficacy of the treatment was evaluated through the monitoring of the survival time of the mice bearing intracranial B16F10 tumor. The STCs+CLX-Lipo@Gel treatment exhibited the most extended survival duration, with a median survival time of 16 days, in contrast to 11.5 days for the STCs+CLX-Lipo group and 9 days for the PBS group (Figure 9B and C).

**Figure 9.**
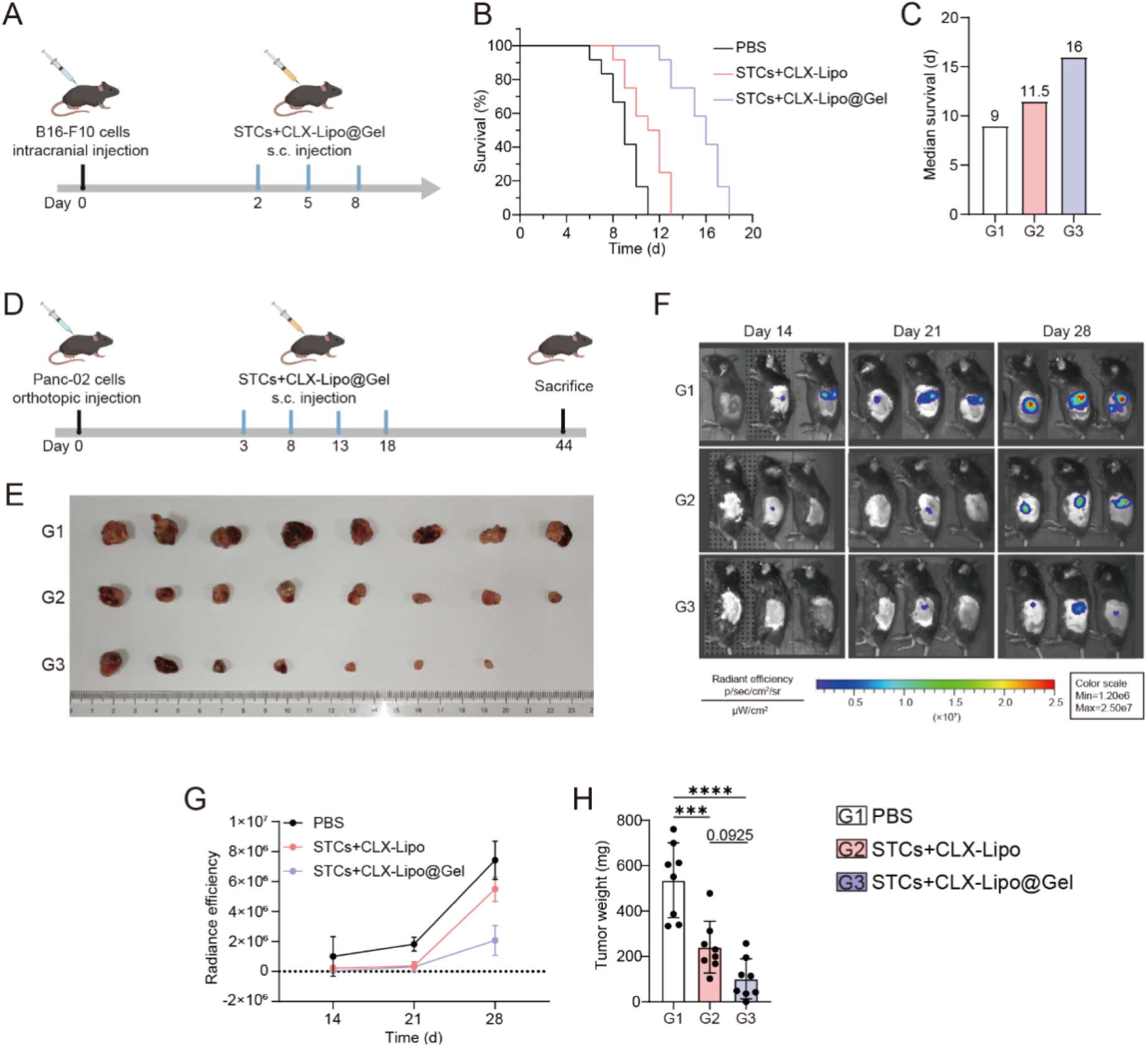
Antitumor effect of STCs+CLX-Lipo@Gel in various tumor models. (A) Schedule of various treatments in melanoma brain metastasis model. Created with BioRender.com (https://BioRender.com/ttybc4v). (B) Survival rate, and (C) median survival of mice during the treatment period (n=12). (D) Schedule of various treatments in an orthotopic pancreatic cancer model. Created with BioRender.com (https://BioRender.com/ttybc4v). (E) Tumor image of pancreatic tumors after different treatments (n=8). (F) Representative IVIS images of pancreatic tumors at different time points after different treatments and (G) semi-quantitative analysis of bioluminescence (n=3). (H) Pancreatic tumor weight of different groups (n=8). Statistical significance was assessed by one-way ANOVA and Tukey multiple comparisons tests. Data were presented as mean ± SD. Statistical significance was determined as ***p < 0.001, ****p < 0.0001.

To further explore the general applicability of the STC-based live vaccine, we conducted a treatment in a typical “cold” tumor model, pancreatic ductal adenocarcinoma (PDAC). In the orthotopic PDAC model, the mice were vaccinated according to the scheme in Figure 9D. The treatment with STCs+CLX-Lipo@Gel showed a significant deceleration in tumor growth (Figure 9E–G), with an average tumor weight of merely 101 mg (Figure 9H). These data underscored the potential of the STCs+CLX-Lipo@Gel as a promising therapeutic strategy, not only in exerting a distant effect on brain metastasis but also in addressing formidable malignancies such as PDAC.

In addition, we designed the study to functionally dissect the relative contribution of senescence-specific neoantigens versus shared tumor antigens. Because both LLC and B16-F10 are syngeneic on the C57BL/6 background, an unmatched senescent-cell vaccine excludes allogeneic rejection while differing from the matched vaccine only in its antigen repertoire. In mice bearing LLC tumors, the matched vaccine (LLC-CLX-Lipo@Gel) reduced tumor burden by ∼90% versus PBS, whereas the unmatched vaccine (B16F10-CLX-Lipo@Gel) achieved only 40% (Figure S9). This difference demonstrates that antitumor immunity elicited by the STC vaccine is directed against tumor antigens shared between senescent cells and their parental tumor cells, rather than depending on senescence-specific neoantigens. However, because this experiment did not include a cell-free arm under identical conditions, we cannot formally partition the residual activity, and a contribution from tumor antigens incidentally shared between two C57BL/6 tumor lines cannot be excluded.

### 2.10 Biosafety evaluation of STCs+CLX-Lipo@Gel

During the treatment period, a slight increase in body weight was observed in all groups of mice, which initially indicates good biosafety of the formulations (Figure S10A). There were no significant differences in organ coefficients among the groups (Figure S10B), and H&E staining results indicated no significant pathological changes in the main organs in all groups (Figure S10C). Moreover, upon subcutaneous injection, STCs exhibited no tumorigenic capacity (Figure S10D). These experimental results suggested that STCs+CLX-Lipo@Gel had good biosafety.

In vivo degradation study showed that the chitosan hydrogel began to degrade around day 3, degraded progressively thereafter, and was completely degraded by day 28 (Figure S11A). It suggested the exposure of the local niche to live STCs and their secretome is temporally bounded (∼4 weeks) rather than indefinite, which substantially limits the risk of chronic inflammation or pro-tumorigenic tissue remodeling. Furthermore, H&E staining revealed no abnormal immune-cell infiltration or tissue damage in the surrounding skin at any time point, indicating local biosafety throughout the entire residence window (Figure S11B).

## 3 Discussion

Tumor vaccines represent a promising therapeutic strategy. Whole tumor cell vaccines, containing the complete spectrum of tumor antigens, can elicit robust local innate immune responses and adaptive anti-tumor responses, garnering significant interest from researchers. STCs, with prolonged activity but without proliferation capacity, possess the potential to gradually release tumor antigens and immunostimulatory cytokines, making them a promising candidate as live vaccines. SASP plays a dual role in cancer progression (Faget et al., 2019; Takasugi et al., 2023). In our study, we proposed an SASP regulation method, in which a negative factor PGE2 was inhibited using CLX-Lipo while SASP action was extended for improving anticancer immunity. It suggested that analysis of SASP components and the use of advanced technologies for comprehensive immune regulation holds broad application prospects.

The hydrogel matrix prolonged the action of STCs, and the co-encapsulated CLX-Lipo promoted the recruitment and maturation of DCs by inhibiting PGE2 production. Our in vivo studies demonstrated that STCs+CLX-Lipo@Gel significantly enhanced NK cell activation, DC maturation, and cDC1 recruitment, culminating in a marked activation of CD8^+^ T cells in the tumor microenvironment. Notably, STCs+CLX-Lipo@Gel extended the survival of mice with melanoma brain metastases and suppressed tumor growth in mice with orthotopic pancreatic tumors, underscoring its potential as a general immunotherapeutic strategy.

Notably, STCs outperformed the died STCs, suggesting the live metabolic state is crucial. It may contribute through two complementary mechanisms. (1) Sustained adjuvant/secretory function: the SASP is an actively regulated, translationally controlled secretory program (Chien et al., 2011; Herranz et al., 2015). Live STCs therefore continually replenish chemokines and cytokines and provide ongoing paracrine stimulation to immature immune cells within the gel depot, whereas a freeze/thaw lysate delivers only a single, finite bolus of pre-formed mediators and DAMPs. A similar advantage was also observed in a killed but metabolically active bacterial vaccine that highly elicits effector T-cell responses and benefits from the potency of live vaccines and the safety of non-proliferation (Brockstedt et al., 2005). (2) Prolonged in situ antigen release: sustained release from an in-situ scaffold outperforms bolus antigen delivery (Kim et al., 2015). In our system, live STCs persist within the gel and release antigens under continuing innate activation, consistent with the observation that senescent cancer cell vaccines outperform immunogenic-cell-death vaccines (Marin et al., 2023a). Regarding the duration, our degradation study shows the hydrogel matrix began degrading at day 3 and was fully degraded by day 28 (Figure S11A), defining the upper bound of the in vivo secretory phase at approximately 4 weeks.

An important question is whether the efficacy of the STC vaccine depends on senescence-specific neoantigens or on the tumor-antigen repertoire shared with parental tumor cells. Our matched-versus-mismatched experiment indicates the latter: in mice bearing LLC tumors, replacing the matched senescent-cell vaccine (LLC-CLX-Lipo@Gel) with a syngeneic but antigen-mismatched senescent-cell vaccine (B16F10-CLX-Lipo@Gel) reduced the tumor inhibition rate from approximately 90% to 40%. The residual activity of the mismatched vaccine is consistent with the antigen-independent, SASP-driven innate component of the platform, as the cell-free components of the formulation (CS@Gel and CLX-Lipo alone) do not restrain tumor growth (Figure 7E–F), whereas senescent tumor cells directly activate NK cells and restore the NK–DC axis (Figures 1E–G and 6). Importantly, this does not diminish the role of the senescence state. Senescence remodels the cell-surface proteome and enhances the antigen-presenting machinery (Chen et al., 2023), and senescent melanocytes upregulate MHC class II and activate T-cell proliferation (van Tuyn et al., 2017); senescent cancer cells are also more efficient than cells undergoing immunogenic cell death at activating DCs and antigen-specific CD8⁺ T cells (Marin et al., 2023a). Consistent with this, live STCs outperform freeze/thaw-killed STCs in our study (Figure 2). We therefore propose that the senescence program contributes to vaccine efficacy by rendering the shared tumor-antigen repertoire more immunologically visible and by providing sustained innate adjuvanticity, rather than by generating a private antigen set. From a translational standpoint this is favorable: the vaccine engages the proliferating tumor that constitutes the bulk of a patient’s tumor burden, rather than the small senescent fraction.

A limitation of this study is that the results do not definitively indicate whether SASP is detrimental to anti-tumor effects. However, extrapolations from in vivo animal experiments suggest that the modified senescent tumor cells may maximize the potential to activate innate and adaptive immunity. Meanwhile, although our findings demonstrate that the STC vaccine could promote the recruitment and maturation of cDC1s with robust CD8^+^ T cell priming, a definitive causal validation remains to be fully established. Specifically, genetic ablation models (such as Batf3^-/-^mice) or systemic cDC1 depletion strategies were not employed in the current study to evaluate whether the therapeutic efficacy is strictly contingent upon cDC1 activation. Future investigations utilizing lineage-deficient models or targeted depletion will be necessary to dissect the indispensable role of this specific DC subset in sustaining long-term antitumor immunity. Another consideration is the inherent heterogeneity of cellular senescence. Emerging evidence underscores that senescent cells do not constitute a homogeneous population; rather, their phenotypic trajectories, metabolic states, and senescence-associated secretory phenotype profiles vary depending on the cell type of origin, the triggering stimuli, and temporal dynamics. Our study predominantly focused on chemotherapy-induced senescence phenotypes characterized by SA-β-gal positivity and cell-cycle arrest markers. Future investigations employing systematic comparison of senescence induced by different stimuli and single-cell transcriptomics will be valuable to comprehensively dissect the functional diversity across senescent subsets in this therapeutic context.

From a translational perspective, senescent cell-based therapies face distinct practical considerations. An autologous approach preserves patient-specific neoantigens but is constrained by limited biopsy materials and lengthy production cycles. A potential application is for postoperative vaccination using surgical tumor tissues as autologous cell source. Alternatively, an ‘off-the-shelf’ allogeneic platform offers superior scalability, standardized biobanking, and consistent batch quality, though potential allo-reactivity must be managed. Furthermore, the clinical administration of live senescent cells presents stringent regulatory hurdles. Key challenges include verifying irreversible growth arrest to prevent malignant escape, controlling heterogeneous SASP-mediated effects, and establishing standardized GMP bioprocesses. Addressing these manufacturing and biosafety criteria will be essential for successful clinical translation.

## 4 Conclusion

Our study has elucidated the promising role of senescent tumor cells, with their inherent high immunogenicity and adjuvanticity, in the development of innovative tumor antigen-based vaccines. Through strategic pharmacological interventions targeting the SASP, particularly the inhibition of PGE2 with CLX, we have effectively restored the function of the NK-DC axis. In vivo experiments demonstrated that the combination of STCs with CLX-Lipo in a chitosan hydrogel matrix effectively activated both innate and adaptive immunity, enhancing the infiltration of immune cells within tumors and reconstructing the tumor immune microenvironment. This strategic modulation was pivotal in amplifying the immune-activating properties of WTC vaccines, thereby contributing to the prevention and suppression of melanoma initiation and progression, and achieving remarkable therapeutic efficacy within a cold tumor model of PDAC. Our study demonstrated that employing senescent tumor cells as a basis for tumor vaccine immunotherapy significantly improved the anti-tumor efficacy of these vaccines, providing novel insights for the development and clinical application of tumor vaccines.

## 5 Material and methods

### 5.1 Materials

Doxorubicin was purchased from Macklin Biochemical Technology Co., Ltd. (Shanghai, China). Chitosan (Cat. No. 448877) was purchased from Sigma-Aldrich Co., Ltd. (St. Louis, USA). PGE2 was purchased from AbMole Bioscience Co., Ltd. (Shanghai, China). PC-98T, cholesterol, and DSPE-MPEG2000 were obtained from AVT Pharmaceutical Tech Co., Ltd. (Shanghai, China). Total RNA extraction reagent, SYBR green master mix, and CellTracker green CMFDA (5-chloromethyl fluorescein diacetate) were purchased from Yeasen Biotechnology Co., Ltd. (Shanghai, China). Murine granulocyte-macrophage colony-stimulating factor (GM-CSF) and murine interleukin 4 (IL-4) were purchased from Novus Biologicals (Centennial, USA). Senescence β-galactosidase staining kit, red blood cell lysis buffer, BCA protein assay kit, crystal violet staining solution, and second antibodies were supplied from Beyotime Biotech Inc (Haimen, China). Mouse IL-6, CXCL1/KC and TNF-α ELISA Kit were purchased from Multi Sciences (Lianke) Biotech Co., Ltd. (Hangzhou, China). Meloxicam, Celecoxib, cell membrane fluorescence probe DiR, CCK-8 kit, D-luciferin potassium, fetal bovine serum (FBS), and Dulbecco’s Modified Eagle’s Medium (DMEM) cell culture medium were purchased from Meilun Biotechnology Co., Ltd. (Dalian, China). COX2 (D5H5) XP® Rabbit mAb was purchased from Cell Signaling Technology. β-Actin Rabbit mAb (AC026) was purchased from ABclonal Technology Co., Ltd. (Wuhan, China). Antibodies used for flow cytometry were purchased from BioLegend (San Diego, USA) and BD Biosciences (San Jose,,USA).

### 5.2 Animals

Male C57BL/6 mice (RRID:MGI:2159769) aged 6-8 weeks, weighing 18-22 g were obtained from the Shanghai Laboratory Animal Center Co., Ltd. (Shanghai, China), and maintained in an SPF-certified facility with access to sterilized food pellets and distilled water, under a 12-hour light/dark cycle. All the animal experimental procedures were approved by the Institutional Animal Care and Use Committee (IACUC), Shanghai Institute of Materia Medica, Chinese Academy of Sciences (IACUC: 2023-07-HYZ-145).

### 5.3 Inclusion and Exclusion Criteria for Animal Studies

To ensure consistency and minimize bias in the animal models, the following criteria were applied:

Inclusion criteria: Only healthy male C57BL/6 mice aged 6–8 weeks, weighing 18–22 g, and confirmed free from specific pathogens (as per supplier certification) were included. Mice were acclimatized for at least 7 days prior to experiments, with normal activity levels and no visible signs of diseases.

Exclusion criteria: Mice exhibiting any abnormalities during acclimatization (e.g., weight loss >10%, lethargy, eating disorders, respiratory distress, or external injuries) were excluded. Tumor-bearing mice were immediately euthanized and excluded from the study if they met any of the following criteria: (1) tumor volume >2000 mm³; (2) any single tumor dimension (length or width) >20 mm. Additionally, mice showing severe distress (e.g., >20% weight loss, immobility, or ulcerated tumors) were euthanized per IACUC guidelines.

### 5.4 Attrition Management

Attrition (loss of animals due to death, euthanasia, or technical issues) was monitored throughout all experimental phases to ensure data integrity:

In survival studies (e.g., melanoma brain metastases model in section 5.14.3), any animal death was recorded daily, with cause documented (e.g., tumor burden, procedural complications, or unrelated morbidity). Animals that died or required early euthanasia were included in survival analyses (e.g., Kaplan-Meier curves) until the point of exit, but excluded from endpoint assays (e.g., tumor weight or immune cell analysis) if tissue was unrecoverable. Attrition rates were minimized by adhering to IACUC guidelines.

### 5.5 Blinding

Treatment administration: All injections were prepared by an independent researcher using coded syringes. Group assignments were concealed from personnel conducting injections and measurements until data analysis.

Tumor assessment: Tumor volume measurements were performed by technicians blinded to treatment groups using anonymized animal IDs.

Survival studies: Endpoint adjudication (death/euthanasia) followed predefined criteria reviewed by two independent blinded investigators.

### 5.6 Power Analysis

Sample size was based on estimations by power analysis using G*Power (RRID:SCR_013726) with a level of significance of 0.05 and a power of 0.85.

### 5.7 Cell culture

The mouse melanoma B16-F10 cell line (RRID:CVCL_0159) was purchased from Meilun Biotechnology Co., Ltd. (Dalian, China). The natural killer cell line NK-92 (RRID: CVCL_2142) and NK-92 cell-specific culture medium were purchased from Wuhan Pronucleus Life Science and Technology Co., Ltd. (Wuhan, China). PANC-02/luci cell line (RRID: CVCL_A8QL) was kindly gifted by Professor Chen Jiang (Fudan University). Lewis Lung Carcinoma cells (LLC) were purchased from the National Collection of Authenticated Cell Cultures, Chinese Academy of Sciences. All cell lines were authenticated by STR profiling tested negative for mycoplasma contamination.

The B16-F10 cells and LLC were cultured in DMEM (MA0212, Meilunbio) with 10% FBS (PWL001, Meilunbio), 100 µg/mL of streptomycin and 100 U/mL of penicillin. The NK-92 cells were cultured in an NK-92 cell-specific culture medium (ProCell, China). All the cells were cultured in a humidified incubator set at 37°C and 5% CO_2_.

### 5.8 Senescence induction and characterization

Senescent B16-F10 cells (STCs) and Senescent LLC cells were induced by treatment with 200 nM doxorubicin for 2–3 days, followed by cultivation for 5–7 days.

#### 5.8.1 β-galactosidase staining assay

β-galactosidase staining assay was conducted using a senescence β-galactosidase staining kit (Beyotime, C0602) according to the manufacturer’s instructions. After incubation, the cells were fixed using β-galactosidase staining fixation solution for 15 min, washed with PBS 3 times, and stained by β-galactosidase staining working solution at 37°C for 12 h. Then, cells were washed with PBS and visualized by Olympus IX73 inverted microscope.

#### 5.8.2 Gene expression analysis by RT-qPCR

Cell total RNA was extracted according to the protocol provided by the total RNA extraction reagent (Yeasen, 10606ES). The concentration of total RNA in each sample was determined using a microvolume spectrophotometer and subsequently equalized. Reverse transcription was carried out following the instructions of the reverse transcription kit (Yeasen, 11151ES10). Quantitative real-time PCR was conducted using SYBR green master mix (Yeasen, 11201ES08) in the Bio-Rad CFX384 Touch Real-

Time PCR Detection System. β-Actin was used as an endogenous control to normalize the relative expression of all target gene mRNAs. The primer sequences are listed in the Table S3.

#### 5.8.3 ELISA for measurement of IL-6, CXCL1, and TNF-α

STCs were induced as indicated, and the supernatant was collected and diluted 5-fold with PBS. The concentrations of cytokines were determined according to the instructions provided by the mouse IL-6 ELISA kit (LIANKEBIO, EK206) and mouse CXCL1/KC ELISA kit (LIANKEBIO, EK296).

Senescent B16F10 cells (2×10^5^ cells/well) were seeded into 12-well plates and cultured for 48 h in medium containing either PBS or celecoxib (5 μM). After incubation, cell culture supernatants were harvested, and the concentrations of IL-6 and TNF-α (LIANKEBIO, EK282) were quantified according to the manufacturer’s instructions.

### 5.9 In vitro study of STCs

#### 5.9.1 Bone marrow-derived dendritic cells (BMDC) generation

Bone marrow cells were obtained from the femurs and tibias of the C57BL/6 mice. The cells were harvested through centrifugation at 150×g for 5 min and subsequently cultured in RPMI1640 supplemented with 20 ng/ml GM-CSF and 10 ng/ml IL-4. Partial medium replacement was performed every 3 days. On days 6–8, non-adherent and loosely adherent immature dendritic cells were collected.

#### 5.9.2 Assessment of DCs maturation

BMDCs (4×10^5^ cells/well) were cocultured with untreated STCs (2×10^5^ cells/well) for 48 h. After incubation, DCs were collected and the expression of maturity markers was analyzed using flow cytometry.

Meanwhile, the culture medium was collected to measure ATP and HMGB1 levels. ATP content determination was performed following the instructions of the enhanced ATP assay kit (Meilunbio, MA0440). The standard and sample were added to each well, and the relative light units (RLU) were measured using BioTek Synergy H1 Multi-Mode Microplate Reader. The determination of HMGB1 content was conducted following the instructions provided in the Mouse High Mobility Group Box 1 (HMGB1) ELISA kit.

#### 5.9.3 Assessment of T cells and NK cells activation

The splenic lymphocytes were isolated from healthy C57BL/6 mice according to the protocol provided by the mouse lymphocyte separation medium (DAKEWE, 7211011). The cells (2×10^6^ cells/well) were cocultured with untreated and senescent B16-F10 cells for 48 h. Then, the cells were collected to assess the activation of T cells and NK cells using flow cytometry. At the same time, the supernatant from the cell culture was collected and the level of perforin and granzyme B was quantified using the mouse perforin ELISA kit and mouse granzyme B ELISA kit respectively, following the provided instructions.

#### 5.9.4 In vitro DCs recruitment

B16-F10 cells were seeded in 12 well plates and induced senescence as indicated. After celecoxib treatment for 48 h, cells were harvested and the expression of COX2 was assessed using western blot analysis. At the same time, the supernatant from the cell culture was collected and the level of PGE2 was quantified using the Mouse Prostaglandin E2 ELISA Kit, following the provided instructions.

STCs and normal B16-F10 cells were seeded in 12 well plates and treated with celecoxib for 48 hours. Following this, lymphocytes isolated from the spleens of mice were seeded into the wells (2×10^6^/well) and further incubated for an additional 48 hours. Subsequently, BMDCs were seeded into the upper chambers of Transwell inserts with a pore size of 8 μm (2×10^5^/well), where they were cocultured with lymphocytes and tumor cells for 48 h. After the completion of the incubation period, the chambers were stained with crystal violet and examined using bright-field microscopy for both counting and photography purposes.

To verify the specificity of the COX-2/PGE2 pathway, senescent B16F10 cells were seeded into 12 well plates (2×10^5^ cells/well) and cultured for 48 h in medium containing different treatments. The experimental groups were designated as follows: celecoxib (5 μM), meloxicam (5 μM), celecoxib (5 μM) + PGE2 (100 ng/mL), meloxicam (5 μM) + PGE2 (100 ng/mL), and a vehicle control (PBS). Following incubation, the medium was replaced with fresh drug-containing medium, and NK92 cells (2×10^5^ cells/well) were added for 48 h of co-culture. Subsequently, a Transwell assay was performed by BMDCs into the upper chambers (8 μm) at a density of 2×10^5^ cells/well. After an additional 48 h of incubation, the inserts were removed, and the migrated BMDCs were stained with crystal violet and examined using bright-field microscopy for photography purposes.

### 5.10 In vivo immunotherapy effect of STCs

Fifteen male C57BL/6 mice received a subcutaneous injection of 4×10^5^ B16-F10 cells in 100 μL PBS on Day 0 and were divided into 3 groups (n=5) on Day 4 randomly. Mice were injected with PBS, Died STCs (STCs dealt with freeze-thaw 3 times), and STCs with 2×10^5^ cells every 2 days for 4 times. The tumor volume was measured every other day. On Day 18, the tumors were excised completely for subsequent examination. On Day 34, mice were sacrificed, and their tumors and spleens were harvested for subsequent analysis.

To analyze the immune cell population in tumors, tumor tissues were dissected into fragments and enzymatically digested using collagenase Ⅳ (1 mg/mL), hyaluronidase (1 mg/mL), and DNase (0.1 mg/mL) at 37 °C for 60 min. The cell suspension was filtered through a 70 μm sieve mesh and stained with primary antibodies of anti-CD45-APC-Cy7 (BD Biosciences Cat# 561586, RRID:AB_10896305), anti-CD11c-PE-Cy7 (BioLegend Cat# 117318, RRID:AB_493568), and anti-MHCII-PE (BD Biosciences Cat# 557000, RRID:AB_396546) for DCs evaluation via flow cytometric analysis. Additionally, to appraise the infiltration of antitumor T lymphocytes, the cell suspension was stained with primary antibodies of anti-CD45-FITC (BioLegend Cat# 372208, RRID:AB_2687032) (BioLegend Cat# 372208, RRID:AB_2687032), anti-CD3-PerCP (BioLegend Cat# 100287, RRID:AB_3683123), anti-CD4-BV605 (BioLegend Cat# 116027, RRID:AB_2800581), anti-CD8-APC (BioLegend Cat# 100712, RRID:AB_312751), anti-IFNγ-PE (BioLegend Cat# 505807, RRID:AB_315401) and anti-Ki67-BV421 (BioLegend Cat# 652411, RRID:AB_2562663), followed by flow cytometric analysis.

To analyze the immune cell population in the spleen, spleens (n=3) were collected and ground to prepare cell suspension. This suspension was then stained with a suite of primary antibodies: anti-CD45-APC-Cy7 (BD Biosciences Cat# 561586, RRID:AB_10896305), anti-NK1.1-APC (BioLegend Cat# 156505, RRID:AB_2876525), anti-CD3-BB700 (BD Biosciences Cat# 566494, RRID:AB_2744393), anti-CD49b-FITC (BioLegend Cat# 103503, RRID:AB_313026), and anti-Granzyme B-PE (BioLegend Cat# 372208, RRID:AB_2687032), for the assessment of NK cells. Concurrently, a suite of primary antibodies: anti-CD45-FITC (BioLegend Cat# 372208, RRID:AB_2687032) (BioLegend Cat# 372208, RRID:AB_2687032), anti-CD3-BB700 (BD Biosciences Cat# 566494, RRID:AB_2744393), anti-CD4-BV605 (BioLegend Cat# 116027, RRID:AB_2800581), anti-CD8-PE (BioLegend Cat# 100708, RRID:AB_312747), anti-CD44-BB515 (BD Biosciences Cat# 564587, RRID:AB_2738855), and anti-CD62L-APC (BD Biosciences Cat# 553152, RRID:AB_398533), was employed for the evaluation of effective memory T cells. These preparations were subjected to flow cytometric analysis to elucidate their immunophenotypes.

### 5.11 Preparation and characterization of CLX-Lipo

The preparation of celecoxib-loaded liposomes (CLX-Lipo) via the thin-film hydration method involved PC-98T, cholesterol, DSPE-MPEG2000, and celecoxib at the mass ratio of 45/5/2/1, respectively. These components were dissolved in the mixed solvent of chloroform/methanol (2/1, v/v). The solution was subjected to rotary evaporation under a water bath at 42°C until the uniformly dispersed film was formed. Subsequently, it was hydrated with water and sonicated using a 270 W probe for 15 min. The free drug was removed using a G50 Sephadex gel column.

The size and zeta potential of CLX-Lipo were measured by the Zeta Sizer Nanoparticle Analyzer (Malvern, UK). Visualization of CLX-Lipo was achieved by negative staining with 1% uranyl acetate followed by examination using transmission electron microscopy. The encapsulation efficiency and drug-loading capacity were assessed via high-performance liquid chromatography (HPLC) (1260 Infinity, Agilent Technologies, Palo Alto, USA), and calculated using the following formula:

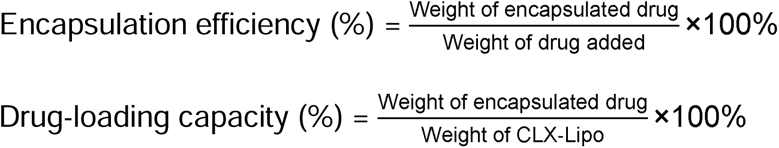

### 5.12 Preparation and characterization of STCs+CLX-Lipo@Gel

The chitosan hydrogel was prepared according to the previously described method. The chitosan solution (3.33%, w/v) was prepared by dissolving chitosan in 0.1 M HCl and sterilized by ultraviolet-light radiation.

The phosphate buffer solution (PB buffer) at pH 8 was prepared by dissolving Na_2_HPO_4_ and NaH_2_PO_4_ in a ratio of 1/0.047 (w/w). The gelling agent was prepared by dissolving NaHCO_3_ in the PB buffer at a molar ratio of 4:5 (NaHCO3 to PB buffer). The gelling agent was sterilized by filtration through 0.2 μm filters.

The formulation of the hydrogel incorporating STCs and CLX-Lipo (denoted as STCs+CLX-Lipo@Gel) was achieved via a sequential two-step procedure. Initially, the chitosan solution was amalgamated with the gelling agent (containing PB buffer and NaHCO_3_ at double concentration) through a meticulous mixing process executed 15 times via a Luer connector. The entire mixture was then transferred into a single syringe. Subsequently, a syringe containing cell suspension with CLX-Lipo was connected via a

Luer connector, and the mixing process was repeated an additional 15 times. The resultant hydrogels were prepared to attain a final concentration of 2% (w/v) chitosan, 0.04 M PB buffer, and 0.05 M NaHCO_3_. Control hydrogels were fabricated following the identical protocol without the inclusion of liposomes and cells.

The rheological assessment of chitosan hydrogel was conducted at a constant angular frequency of 1 rad/s and shear strain of 1%. The internal morphology of the chitosan hydrogel was observed using scanning electron microscopy. Briefly, the chitosan hydrogel was rapidly frozen in liquid nitrogen and lyophilized, and at least three random regions were captured to observe its morphology.

### 5.13 Viability assay of STCs in chitosan hydrogel

The harvested STCs (5×10^5^ cells/mL) were suspended in chitosan hydrogels of varying concentrations, 0.05 M, 0.075 M, and 0.1 M, and subsequently incubated within a 12-well culture plate for 14 days. After culture, a live/dead assay was conducted to evaluate the viability of the cells. 500 μL Calcein AM/PI detection solution was added to each well, followed by incubation of the cells at 37°C in the dark for 30 min. After washing with PBS, the cells were observed under a fluorescence microscope (Olympus IX73, Japan), and images were captured to calculate the proportion of viable cells.

### 5.14 In vitro study of STCs+CLX-Lipo@Gel

#### 5.14.1 Maturation of BMDCs

Immature BMDCs were cocultured for 48 h with various treatments, including PBS, CS@Gel, CLX-Lipo (5 μM), STCs, STCs+CLX-Lipo (5 μM) and STCs+CLX-Lipo@Gel (5 μM) for 48 h. The cells were stained with primary antibodies of anti-CD11c-FITC (BioLegend Cat# 117306, RRID:AB_313775), anti-MHCII-PE (BD Biosciences Cat# 557000, RRID:AB_396546), anti-CD11b-AF647 (BioLegend Cat# 101218, RRID:AB_389327), and anti-CD103-BV405 (Proteintech Cat# CL405-65047, RRID:AB_3083818). The cellular subtypes were subsequently analyzed using flow cytometry.

#### 5.14.2 Activation of NK cells

NK-92 cells were plated at a density of 2×10^5^ cells per well in a 12-well plate and cocultured with the aforementioned treatments for 48 h. Subsequently, the cells were harvested, and the expression levels of NKp46 and NKG2D on NK cells were assessed using flow cytometry.

To assess the cytotoxicity of NK cells, NK-92 cells were plated at a density of 2×10^5^ cells per well in a 12-well plate and cocultured with the aforementioned treatments for 48 h. Subsequently, the NK cells were harvested and cocultured with B16-F10 cells for 18 hours. After co-incubation, the cells were gently rinsed with PBS to eliminate the NK cells. Then, 500 μL of CCK-8 assay working solution was added to each well, and the plate was further incubated in a cell culture incubator for 2 h. The optical density (OD) was measured at 450 nm, with normal B16-F10 cells used as the control for determining the cell viability rate.

To evaluate the capacity of NK cells to accumulate DCs, NK-92 cells were plated at a density of 2×10^5^ cells per well in a 12-well plate and cocultured with the aforementioned treatments for 48 h. Thereafter, freshly prepared BMDCs were seeded into the upper chambers of Transwell inserts with a pore size of 8 μm (2×10^5^/well) and co-incubated for another 48 h. After the completion of the incubation period, the chambers were subjected to staining with crystal violet and subsequently analyzed via bright-field microscopy for both counting and photography purposes. The supernatant derived from the coculture was subsequently collected and analyzed for the secretion of CCL5, XCL1, and PGE2 using ELISA kits according to the protocols.

#### 5.14.3 Activation of T cells

Lymphocytes extracted from murine spleens were seeded at a density of 1×10^6^ cells per well in a 12-well plate and cocultured with the aforementioned treatments for 48 h. The cells were then harvested and stained with primary antibodies of anti-CD3-PerCP-Cy5.5 (BioLegend Cat# 100218, RRID:AB_1595492 (BioLegend Cat# 100218, RRID:AB_1595492), anti-CD8-BV421 (BioLegend Cat# 100753, RRID:AB_2562558), anti-Granzyme B-PE (BioLegend Cat# 372208, RRID:AB_2687032), and anti-IFNγ-APC-Cy7 (BioLegend Cat# 505849, RRID:AB_2616697). The cellular subtypes were subsequently analyzed using flow cytometry.

#### 5.14.4 Stimulation of T and NK cell proliferation

Lymphocytes harvested from mouse spleens were stained with CMFDA before being seeded at a density of 1×10^6^ cells per well in a 12-well plate and cocultured with the aforementioned treatments for 48 h. Following this, the cells were harvested and stained with primary antibodies of anti-CD3-PerCP-Cy5.5 (BioLegend Cat# 100218, RRID:AB_1595492 (BioLegend Cat# 100218, RRID:AB_1595492), anti-CD8-BV421 (BioLegend Cat# 100753, RRID:AB_2562558), and anti-NK1.1-APC (BioLegend Cat# 156505, RRID:AB_2876525). The proportion of proliferating cells was analyzed using cytometry.

### 5.15 In vivo retention and uptake of STCs in lymph nodes

Male C57BL/6 mice were divided into 2 groups (n=3). STCs were harvested as described above and incubated with DiR cell membrane fluorescent probe staining solution at 37°C for 30 min. The chitosan hydrogel encapsulating STCs, referred to as STCs@Gel, was according to the previously outlined procedure. Subsequently, STCs suspension and STCs@Gel, which had consistent fluorescence intensity, were subcutaneously injected. The fluorescence intensity was quantified utilizing the IVIS imaging system (Caliper PerkinElmer, Hopkinton, MA, USA) on days 1, 2, 3, 5, 7, and 14, followed by statistical analysis of the obtained data.

### 5.16 In vivo immunization and immunotherapy study of STCs+CLX-Lipo@Gel

#### 5.16.1 Prophylactic study in melanoma

Thirty male C57BL/6 mice were divided into 6 groups (n=5) as delineated below: PBS, CS@Gel, CLX-Lipo, STCs, STCs+CLX-Lipo, STCs+CLX-Lipo@Gel. The mice underwent immunization with 2×10^5^ STCs, administered subcutaneously, either in the presence or absence of liposomal CLX at a dosage of 2 mg/kg. 7 days after the immunization, the mice were challenged with 1×10^6^ B16-F10 cells administered subcutaneously into their left flank. The tumor volumes were recorded every other day. On the 14th day, the mice were euthanized, and tissue samples were procured for subsequent experimental analysis.

On the 7th day following immunization, 200 μL of blood was collected from the mice’s posterior orbital venous plexus. Following erythrocyte lysis, the cells were stained with primary antibodies of anti-CD3-PerCP-Cy5.5 (BioLegend Cat# 100218, RRID:AB_1595492 (BioLegend Cat# 100218, RRID:AB_1595492), anti-NK1.1-APC (BioLegend Cat# 156505, RRID:AB_2876525), anti-NKG2D-PE-Dazzle594 (BioLegend Cat# 130213, RRID:AB_2728147), and anti-NKp46-BV421 (BioLegend Cat# 137611, RRID:AB_10915472). The cellular subtypes were subsequently analyzed via flow cytometry.

On the 14th day following the mice euthanized, spleens were harvested and processed to create a cell suspension through grinding. The cells were stained with primary antibodies of anti-CD3-PerCP-Cy5.5 (BioLegend Cat# 100218, RRID:AB_1595492 (BioLegend Cat# 100218, RRID:AB_1595492, anti-CD4-FITC (BD Biosciences Cat# 553079, RRID:AB_394609), anti-CD8-BV421 (BioLegend Cat# 100753, RRID:AB_2562558), anti-CD44-PE (BD Biosciences Cat# 553134, RRID:AB_394649), and anti-CD62L-APC (BD Biosciences Cat# 553152, RRID:AB_398533). The cellular subtypes were subsequently analyzed via flow cytometry.

#### 5.16.2 Immunotherapy study in melanoma

Twenty-five male C57BL/6 mice were administered a subcutaneous injection of 4×10^5^ B16-F10 cells suspended in 100 μL PBS on Day 0. These mice were randomly divided into 5 groups (n=5) as follows: PBS, CLX-Lipo, STCs, STCs+CLX-Lipo, STCs+CLX-Lipo@Gel. The dosages administered were consistent with the aforementioned specifications on the 11th day. The tumor volumes were recorded every other day. On the 20th day, the mice were euthanized, and tissue samples were harvested for subsequent experimental analysis.

To analyze the DCs population in lymph nodes, inguinal lymph nodes were collected and grinded to obtain the single-cell suspension. The cells were stained with primary antibodies of anti-CD45-FITC (BioLegend Cat# 372208, RRID:AB_2687032), anti-CD11c-PE-Cy7 (BioLegend Cat# 117318, RRID:AB_493568), anti-MHCII-PE (BD Biosciences Cat# 557000, RRID:AB_396546), anti-CD11b-AF647 (BioLegend Cat# 101218, RRID:AB_389327), and anti-CD103-BV405 (Proteintech Cat# CL405-65047, RRID:AB_3083818), to facilitate the assessment of DCs via flow cytometric analysis.

To analyze the immune cell population in tumors, the cell suspension was obtained as mentioned above. The cell suspension was stained with primary antibodies of anti-CD45-FITC (BioLegend Cat# 372208, RRID:AB_2687032), anti-CD3-PerCP-Cy5.5 (BioLegend Cat# 100218, RRID:AB_1595492, anti-NK1.1-APC (BioLegend Cat# 156505, RRID:AB_2876525), anti-NKG2D-PE-Dazzle594 (BioLegend Cat# 130213, RRID:AB_2728147), and anti-NKp46-BV421 (BioLegend Cat# 137611, RRID:AB_10915472) for NK cells evaluation via flow cytometric analysis. Additionally, to evaluate the infiltration of antitumor T lymphocytes, the cell suspension was stained with primary antibodies of anti-CD45-FITC (BioLegend Cat# 372208, RRID:AB_2687032), anti-CD3-PerCP (BioLegend Cat# 100287, RRID:AB_3683123), anti-CD8-BV421 (BioLegend Cat# 100753, RRID:AB_2562558), anti-IFNγ-PE-Cy7 (BioLegend Cat# 505825, RRID:AB_1595591), anti-Granzyme B-PE (BioLegend Cat# 372208, RRID:AB_2687032), and anti-Ki67-APC (BioLegend Cat# 652405, RRID:AB_2561929), followed by flow cytometric analysis.

#### 5.16.3 In Vivo Validation of Antigen Specificity of STCs vaccine

Healthy C57BL/6 mice were subcutaneously inoculated with LLC cells (5×10^5^ cells/mouse) into the right flank to establish a subcutaneous tumor model, with the day of inoculation designated as Day 0. On Day 11, tumor-bearing mice were randomized into three groups (n = 5) based on tumor volumes and administered a single subcutaneous injection of PBS, LLC-CLX-Lipo@Gel (Senescent LLC cells, 2×10^5^ cells/mouse), or B16F10-CLX-Lipo@Gel (Senescent B16-F10 cells, 2×10^5^ cells/mouse). Tumor volume (length and width) and mouse body weights were recorded every other day. When the tumor volume reached approximately 2000 mm^3^, the mice were euthanized, and the tumors were excised and weighed.

#### 5.16.4 Therapeutic efficacy in melanoma brain metastases model

A mouse model for melanoma brain metastases was established through intracranial injections of 5×10^3^ B16-F10 cells in 5 μL of PBS into the striatum, located 2 mm right lateral to the bregma, of male C57BL/6 mice. These mice were randomly divided into 3 groups as follows: PBS, STCs+CLX-Lipo, STCs+CLX-Lipo@Gel. Each mouse received subcutaneous immunizations with 5×10^5^ STCs and liposomal CLX at a dosage of 2 mg/kg on days 3, 6, and 9. In the survival study, the survival rate was recorded (n=10).

#### 5.16.5 Therapeutic efficacy in orthotopic pancreatic ductal adenocarcinoma model

Senescent PANC-02 cells were induced by treatment with 200 nM doxorubicin for 2 days, followed by cultivation for 5–7 days. The orthotopic PANC-02 PDAC model was established by injecting 2×10^6^ PANC-02/luci cells suspended in 50 μL matrigel directly into the pancreas of male C57BL/6 mice. The mice were randomly divided into 3 groups as follows: PBS, STCs+CLX-Lipo, STCs+CLX-Lipo@Gel. Each mouse received subcutaneous immunizations with 5×10^5^ STCs and liposomal CLX at a dosage of 2 mg/kg on days 3, 7, 11, and 15. D-luciferin potassium solution was administered intraperitoneally at a dose of 150 mg/kg. Bioluminescence was then detected using the IVIS Spectrum imaging system, 10 minutes after injection, every 7 days. On day 46 post-tumor inoculation, the mice were euthanized, and the tumors were harvested for further analysis.

### 5.17 Biosafety evaluation

The body weight of the mice was measured during the treatment. After the end of the experiment, the hearts, livers, spleens, lungs, and kidneys of the mice were harvested. After being thoroughly washed with PBS and excess liquid absorbed with filter paper, they were weighed and recorded to calculate the organ coefficients.

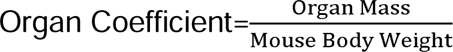

The organ tissues were then immersed in a 4% paraformaldehyde solution for fixation and stored at 4°C. The samples underwent paraffin embedding, tissue sectioning, and H&E staining. Pathological sections were scanned using a pathology slide scanner to assess pathological changes in the organs of mice from different groups, providing a preliminary evaluation of safety.

In the tumorigenesis experiment, 12 male C57BL/6 mice were randomly segregated into three groups, each receiving a subcutaneous injection of 1×10^6^, 2×10^6^, and 5×10^6^ B16-F10 cells suspended in 100 μL PBS respectively, with subsequent monitoring for tumor development.

### 5.18 Hydrogel degradation and biosafety

The hydrogel vaccine was injected subcutaneously into mice, with the injection day designated as Day 0. On Days 1, 3, 7, 14, 21, and 28, the skin tissues surrounding the injection sites were excised and subjected to hematoxylin and eosin (H&E) staining. Concurrently, mice receiving the hydrogel vaccine were photographed at the corresponding time points (Days 1, 3, 7, 14, 21, and 28) to monitor the in vivo degradation profile of the hydrogel depots.

### 5.19 Statistical analysis

All data analysis in this study was performed using GraphPad Prism 9.0 software (RRID:SCR_002798). The data were presented as mean ± standard deviation. For multiple groups (three or more), statistical analysis was performed using one-way analysis of variance (ANOVA); for two groups, a t-test was employed. Statistical significance was determined as *p < 0.05, **p < 0.01, ***p < 0.001, ****p < 0.0001.

## Acknowledgments

We thank the staff members of the Large-scale Protein Preparation System at the National Facility for Protein Science in Shanghai (NFPS), Shanghai Advanced Research Institute, Chinese Academy of Science, for providing technical support and assistance in data collection and analysis. Special thanks are to Prof. Chen Jiang at Fudan University for providing the PANC02/luci cell lines. Some schematic elements were created with BioRender.com.

## Author Contributions

Yuewei Wang: conceptualization, methodology, investigation, formal analysis, data curation, and writing – original draft.

Ante Ou: methodology, investigation, and writing – review and editing. Yanli Luo: investigation and writing – review and editing.

Yanrong Gao: methodology and investigation. Yi Zhang: methodology and investigation.

Linxi Qin: investigation.

Yongzhuo Huang*: conceptualization, methodology, writing – review and editing, supervision, and funding acquisition.

## Availability of data and materials

The dataset supporting the conclusions of this article is included within the article and its SI file.

## Ethics statement

All the animal experimental procedures were approved by the Institutional Animal Care and Use Committee (IACUC), Shanghai Institute of Materia Medica, Chinese Academy of Sciences (IACUC: 2023-07-HYZ-145).

*During the preparation of this work the authors used an AI tool (Deepseek-V4.1-Flash) to improve readability and language of this manuscript. After using this tool, the authors reviewed and edited the content as needed and take full responsibility for the content of the publication*.

## Abbreviations

cDC1: Conventional type 1 dendritic cells
CLX: Celecoxib
DAMPs: Damage-associated molecular patterns
DCs: Dendritic cells
DMEM: Dulbecco’s Modified Eagle’s Medium
FBS: Fetal bovine serum
HMGB1: High mobility group box-1 protein
HPLC: High-performance liquid chromatography
NK: Natural killer
PB: Phosphate buffer
PDAC: Pancreatic ductal adenocarcinoma
PGE2: Prostaglandin E2
SaβG: Senescence-associated β-galactosidase
SASP: Senescence-associated secretory phenotype
STCs: Senescent tumor cells
SEM: Scanning electron microscope
TAAs: Tumor-associated antigens
TSAs: Tumor-specific antigens
TEMs: Effector memory T cells
WB: Western blot
WTC: Whole tumor cell

## Reference

Ahmadi, R., and J.D. de Bruijn. 2008. Biocompatibility and gelation of chitosan-glycerol phosphate hydrogels. J Biomed Mater Res A 86a:824–832.

Antonangeli, F., A. Soriani, B. Ricci, A. Ponzetta, G. Benigni, S. Morrone, G. Bernardini, and A. Santoni. 2016. Natural killer cell recognition of drug-induced senescent multiple myeloma cells. Oncoimmunology 5: e1218105

Antonangeli, F., A. Zingoni, A. Soriani, and A. Santoni. 2019. Senescent cells: Living or dying is a matter of NK cells. J Leukocyte Biol 105:1275–1283.

Assaad, E., M. Maire, and S. Lerouge. 2015. Injectable thermosensitive chitosan hydrogels with controlled gelation kinetics and enhanced mechanical resistance. Carbohyd Polym 130:87–96.

Bluth, M.J., L.C. Zaba, D. Moussai, M. Suárez-Fariñas, H. Kaporis, L. Fan, K.C. Pierson, T.R. White, A. Pitts-Kiefer, J. Fuentes-Duculan, E. Guttman-Yassky, J.G. Krueger, M.A. Lowes, and J.A. Carucci. 2009. Myeloid Dendritic Cells from Human Cutaneous Squamous Cell Carcinoma Are Poor Stimulators of T-Cell Proliferation. J Invest Dermatol 129:2451–2462.

Böttcher, J.P., E. Bonavita, P. Chakravarty, H. Blees, M. Cabeza-Cabrerizo, S. Sammicheli, N.C. Rogers, E. Sahai, S. Zelenay, and C.R.E. Sousa. 2018. NK Cells Stimulate Recruitment of cDC1 into the Tumor Microenvironment Promoting Cancer Immune Control. Cell 172:1022–1037.e14.

Böttcher, J.P., and C.R.E. Sousa. 2018. The Role of Type 1 Conventional Dendritic Cells in Cancer Immunity. Trends in Cancer 4:784–792.

Brockstedt, D.G., K.S. Bahjat, M.A. Giedlin, W. Liu, M. Leong, W. Luckett, Y. Gao, P. Schnupf, D. Kapadia, G. Castro, J.Y. Lim, A. Sampson-Johannes, A.A. Herskovits, A. Stassinopoulos, H.G. Bouwer, J.E. Hearst, D.A. Portnoy, D.N. Cook, and T.W. Dubensky, Jr. 2005. Killed but metabolically active microbes: a new vaccine paradigm for eliciting effector T-cell responses and protective immunity. Nat Med 11:853–860.

Chen, H.A., Y.J. Ho, R. Mezzadra, J.M. Adrover, R. Smolkin, C.Y. Zhu, K. Woess, N. Bernstein, G. Schmitt, L. Fong, W. Luan, A. Wuest, S. Tian, X. Li, C. Broderick, R.C. Hendrickson, M. Egeblad, Z.H. Chen, D. Alonso-Curbelo, and S.W. Lowe. 2023. Senescence Rewires Microenvironment Sensing to Facilitate Antitumor Immunity. Cancer Discovery 13:432–453.

Chien, Y., C. Scuoppo, X. Wang, X. Fang, B. Balgley, J.E. Bolden, P. Premsrirut, W. Luo, A. Chicas, C.S. Lee, S.C. Kogan, and S.W. Lowe. 2011. Control of the senescence-associated secretory phenotype by NF-kappaB promotes senescence and enhances chemosensitivity. Genes Dev 25:2125–2136.

Deng, J.J., W.D. Xu, S.Y. Lei, W.Y. Li, Q.H. Li, K.Q. Li, J.X. Lyu, J.L. Wang, and Z. Wang. 2022. Activated Natural Killer Cells-Dependent Dendritic Cells Recruitment and Maturation by Responsive Nanogels for Targeting Pancreatic Cancer Immunotherapy. Small 18: e2203114

Faget, D.V., Q.H. Ren, and S.A. Stewart. 2019. Unmasking senescence: context-dependent effects of SASP in cancer. Nature Reviews Cancer 19:439–453.

Ferris, S.T., V. Durai, R. Wu, D.J. Theisen, J.P. Ward, M.D. Bern, J.T. Davidson, P. Bagadia, T.T. Liu, C.G. Briseño, L.J. Li, W.E. Gillanders, G.F. Wu, W.M. Yokoyama, T.L. Murphy, R.D. Schreiber, and K.M. Murphy. 2020. cDC1 prime and are licensed by CD4^+^ T cells to induce anti-tumour immunity. Nature 584:624–629.

Gasek, N.S., G.A. Kuchel, J.L. Kirkland, and M. Xu. 2021. Strategies for targeting senescent cells in human disease. Nature Aging 1:870–879.

Gorgoulis, V., P.D. Adams, A. Alimonti, D.C. Bennett, O. Bischof, C. Bishop, J. Campisi, M. Collado, K. Evangelou, G. Ferbeyre, J. Gil, E. Hara, V. Krizhanovsky, D. Jurk, A.B. Maier, M. Narita, L. Niedernhofer, J.F. Passos, P.D. Robbins, C.A. Schmitt, J. Sedivy, K. Vougas, T. Von Zglinicki, D. Zhou, M. Serrano, and M. Demaria. 2019. Cellular Senescence: Defining a Path Forward. Cell 179:813–827.

Hasegawa, T., T. Oka, H.G. Son, V.S. Oliver-García, M. Azin, T.M. Eisenhaure, D.J. Lieb, N. Hacohen, and S. Demehri. 2023. Cytotoxic CD4^+^ T cells eliminate senescent cells by targeting cytomegalovirus antigen. Cell 186:1417–1431.e20.

Herranz, N., S. Gallage, M. Mellone, T. Wuestefeld, S. Klotz, C.J. Hanley, S. Raguz, J.C. Acosta, A.J. Innes, A. Banito, A. Georgilis, A. Montoya, K. Wolter, G. Dharmalingam, P. Faull, T. Carroll, J.P. Martinez-Barbera, P. Cutillas, F. Reisinger, M. Heikenwalder, R.A. Miller, D. Withers, L. Zender, G.J. Thomas, and J. Gil. 2015. mTOR regulates MAPKAPK2 translation to control the senescence-associated secretory phenotype. Nat Cell Biol 17:1205–1217.

Iannello, A., T.W. Thompson, M. Ardolino, S.W. Lowe, and D.H. Raulet. 2013. p53-dependent chemokine production by senescent tumor cells supports NKG2D-dependent tumor elimination by natural killer cells. J Exp Med 210:2057–2069.

Jahani, V., M. Yazdani, A. Badiee, M.R. Jaafari, and L. Arabi. 2023. Liposomal celecoxib combined with dendritic cell therapy enhances antitumor efficacy in melanoma. Journal of Controlled Release 354:453–464.

Kim, J., W.A. Li, Y. Choi, S.A. Lewin, C.S. Verbeke, G. Dranoff, and D.J. Mooney. 2015. Injectable, spontaneously assembling, inorganic scaffolds modulate immune cells in vivo and increase vaccine efficacy. Nat Biotechnol 33:64–72.

Kyrysyuk, O., and K.W. Wucherpfennig. 2023. Designing Cancer Immunotherapies That Engage T Cells and NK Cells. Annu Rev Immunol 41:17–38.

Lee, S., and C.A. Schmitt. 2019. The dynamic nature of senescence in cancer. Nat Cell Biol 21:94–101.

Liu, Y., J. Pagacz, D.J. Wolfgeher, K.D. Bromerg, J.V. Gorman, and S.J. Kron. 2023. Senescent cancer cell vaccines induce cytotoxic T cell responses targeting primary tumors and disseminated tumor cells. J Immunother Cancer 11: e005862

Marin, I., O. Boix, A. Garcia-Garijo, I. Sirois, A. Caballe, E. Zarzuela, I. Ruano, C.S.O. Attolini, N. Prats, J.A. López-Domínguez, M. Kovatcheva, E. Garralda, J. Muñoz, E. Caron, M. Abad, A. Gros, F. Pietrocola, and M. Serrano. 2023a. Cellular Senescence Is Immunogenic and Promotes Antitumor Immunity. Cancer Discovery 13:410–431.

Marin, I., M. Serrano, and F. Pietrocola. 2023b. Recent insights into the crosstalk between senescent cells and CD8 T lymphocytes. npj Aging 9: 8

Meng, Y., E.V. Efimova, K.W. Hamzeh, T.E. Darga, H.J. Mauceri, Y.-X. Fu, S.J. Kron, and R.R. Weichselbaum. 2012. Radiation-inducible Immunotherapy for Cancer: Senescent Tumor Cells as a Cancer Vaccine. Molecular Therapy 20:1046–1055.

Monette, A., C. Ceccaldi, E. Assaad, S. Lerouge, and R. Lapointe. 2016. Chitosan thermogels for local expansion and delivery of tumor-specific T lymphocytes towards enhanced cancer immunotherapies. Biomaterials 75:237–249.

Oliva, I.C.G., G. Schvartsman, and H. Tawbi. 2018. Advances in the systemic treatment of melanoma brain metastases. Ann Oncol 29:1509–1520.

Park, S.S., Y.W. Choi, J.H. Kim, H.S. Kim, and T.J. Park. 2021. Senescent tumor cells: an overlooked adversary in the battle against cancer. Exp Mol Med 53:1834–1841.

Prieto, L.I., I. Sturmlechner, J.J. Goronzy, and D.J. Baker. 2023. Senescent cells as thermostats of antitumor immunity. Sci Transl Med 15: eadg7291

Ruscetti, M., J.P.t. Morris, R. Mezzadra, J. Russell, J. Leibold, P.B. Romesser, J. Simon, A. Kulick, Y.J. Ho, M. Fennell, J. Li, R.J. Norgard, J.E. Wilkinson, D. Alonso-Curbelo, R. Sridharan, D.A. Heller, E. de Stanchina, B.Z. Stanger, C.J. Sherr, and S.W. Lowe. 2020. Senescence-Induced Vascular Remodeling Creates Therapeutic Vulnerabilities in Pancreas Cancer. Cell 181:424–441 e421.

Takasugi, M., Y. Yoshida, E. Hara, and N. Ohtani. 2023. The role of cellular senescence and SASP in tumour microenvironment. Febs J 290:1348–1361.

van Tuyn, J., F. Jaber-Hijazi, D. MacKenzie, J.J. Cole, E. Mann, J.S. Pawlikowski, T.S. Rai, D.M. Nelson, T. McBryan, A. Ivanov, K. Blyth, H. Wu, S. Milling, and P.D. Adams. 2017. Oncogene-Expressing Senescent Melanocytes Up-Regulate MHC Class II, a Candidate Melanoma Suppressor Function. J Invest Dermatol 137:2197–2207.

Von Bergwelt-Baildon, M.S., A. Popov, T. Saric, J. Chemnitz, S. Classen, M.S. Stoffel, F. Fiore, U. Roth, M. Beyer, S. Debey, C. Wickenhauser, F.-G. Hanisch, and J.L. Schultze. 2006. CD25 and indoleamine 2,3-dioxygenase are up-regulated by prostaglandin E2 and expressed by tumor-associated dendritic cells in vivo: additional mechanisms of T-cell inhibition. Blood 108:228–237.

Wculek, S.K., F.J. Cueto, A.M. Mujal, I. Melero, M.F. Krummel, and D. Sancho. 2020. Dendritic cells in cancer immunology and immunotherapy. Nat Rev Immunol 20:7–24.

