## Supplemental Data 1 for "Turning aging cells into a live vaccine: engineered senescent cancer cells with adjuvant celecoxib for immunotherapy"

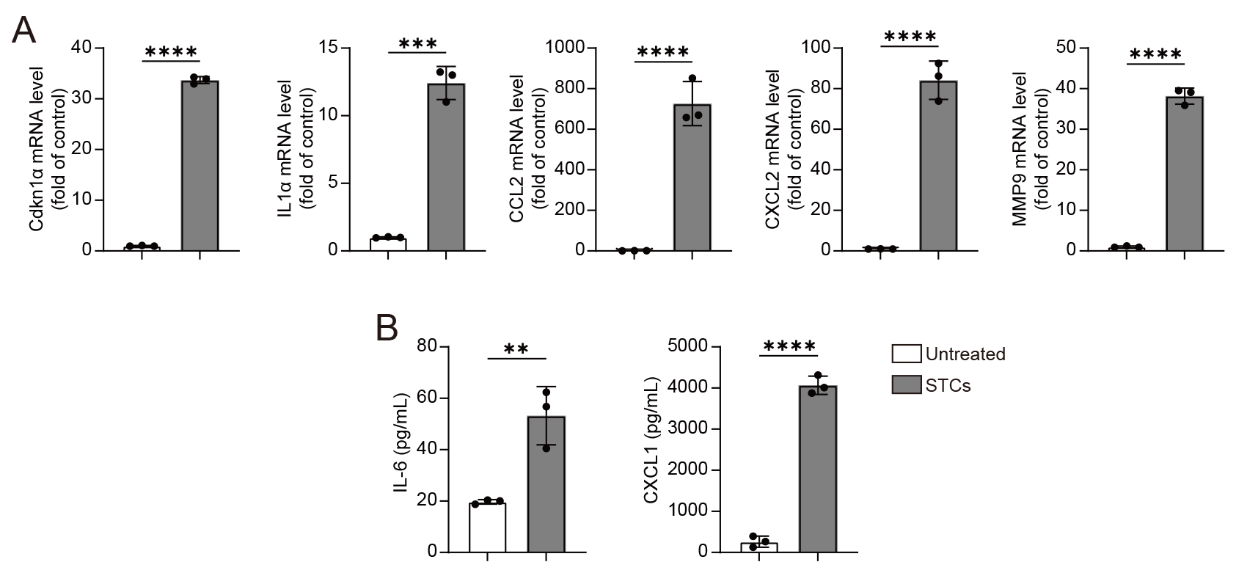

**Figure S1.** Senescent B16-F10 cells characterization. (A) mRNA expression levels of *Cdkn1α*, *IL-1α*, *CCL2*, *CXCL2*, and *MMP9* of senescent B16-F10 cells; (B) The concentrations of IL-6 and CXCL1 in the supernatant of senescent B16-F10 cells. Statistical significance was assessed by unpaired Student’s t-test. Data were presented as mean ± SD. Statistical significance was determined as **p < 0.01, ***p < 0.001, ****p < 0.0001.

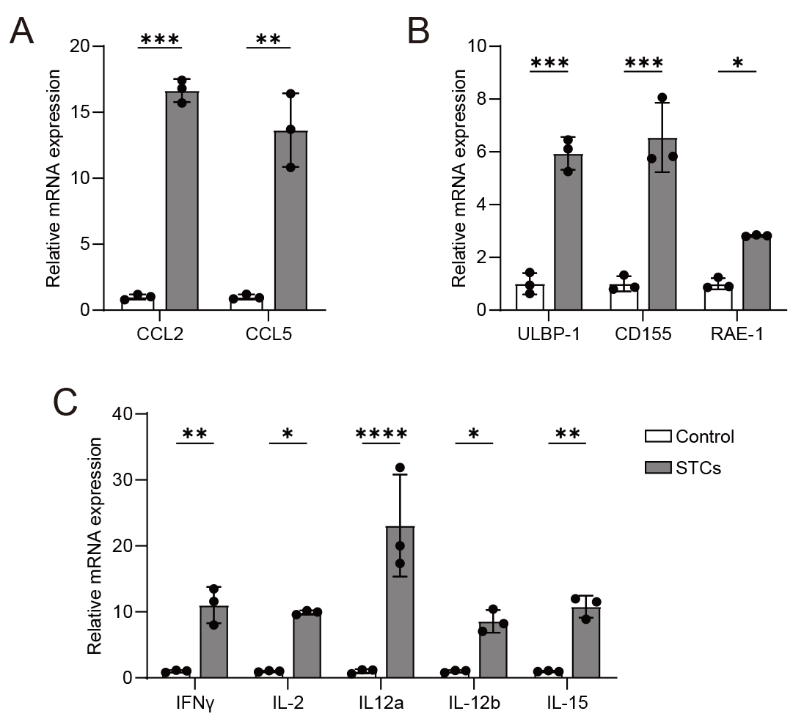

**Figure S2.** Alterations in immune-related cytokines and NK cell-activated receptor ligands at the gene expression level. (A) mRNA expression levels of chemokines CCL2 and CCL5 in STCs; (B) mRNA expression levels of NKG2D ligands ULBP-1 and RAE-1, and DNAM-1 ligand CD155 in STCs; (C) mRNA expression levels of immune-related cytokines IFN-γ, IL-2, IL-12, and IL-15. Statistical significance was assessed by unpaired Student’s t-test. Data were presented as mean ± SD. Statistical significance was determined as *p < 0.05, **p < 0.01, ***p < 0.001, ****p < 0.0001.

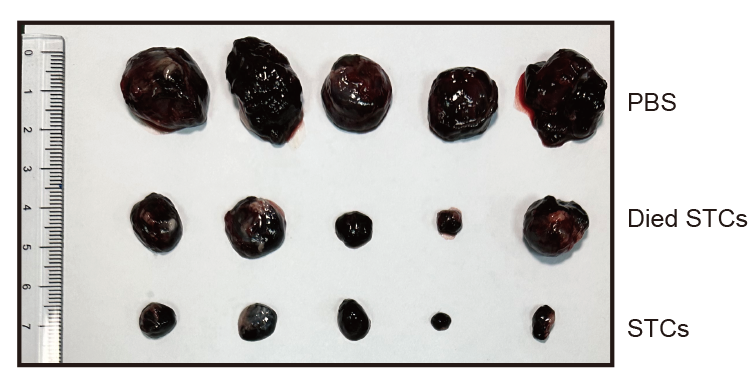

**Figure S3.** Tumor image of the first tumor (n=5).

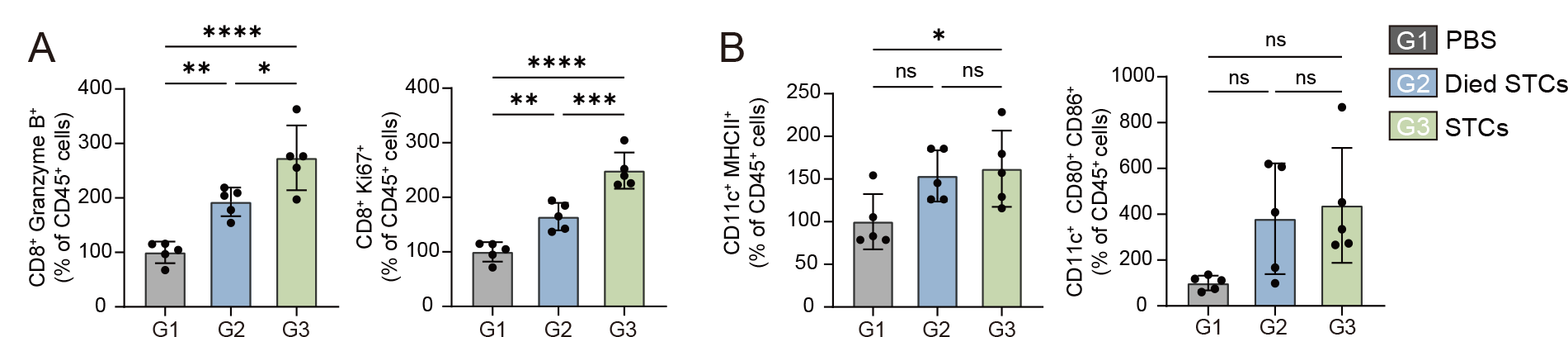

**Figure S4.** The percentage of (A) activated CD8^+^ T cells (CD8^+^ Granzyme B^+^), proliferative CD8^+^ T cells (CD8^+^ Ki67^+^), (B) matured DCs (CD11c^+^ MHCII^+^ and CD11c^+^ CD80^+^ CD86^+^) in tumor tissues after different treatments. Statistical significance was assessed by one-way ANOVA and Tukey multiple comparisons tests. Data were presented as mean ± SD. Statistical significance was determined as ns (not significant, p > 0.05), *p < 0.05, **p < 0.01, ***p < 0.001, ****p < 0.0001.

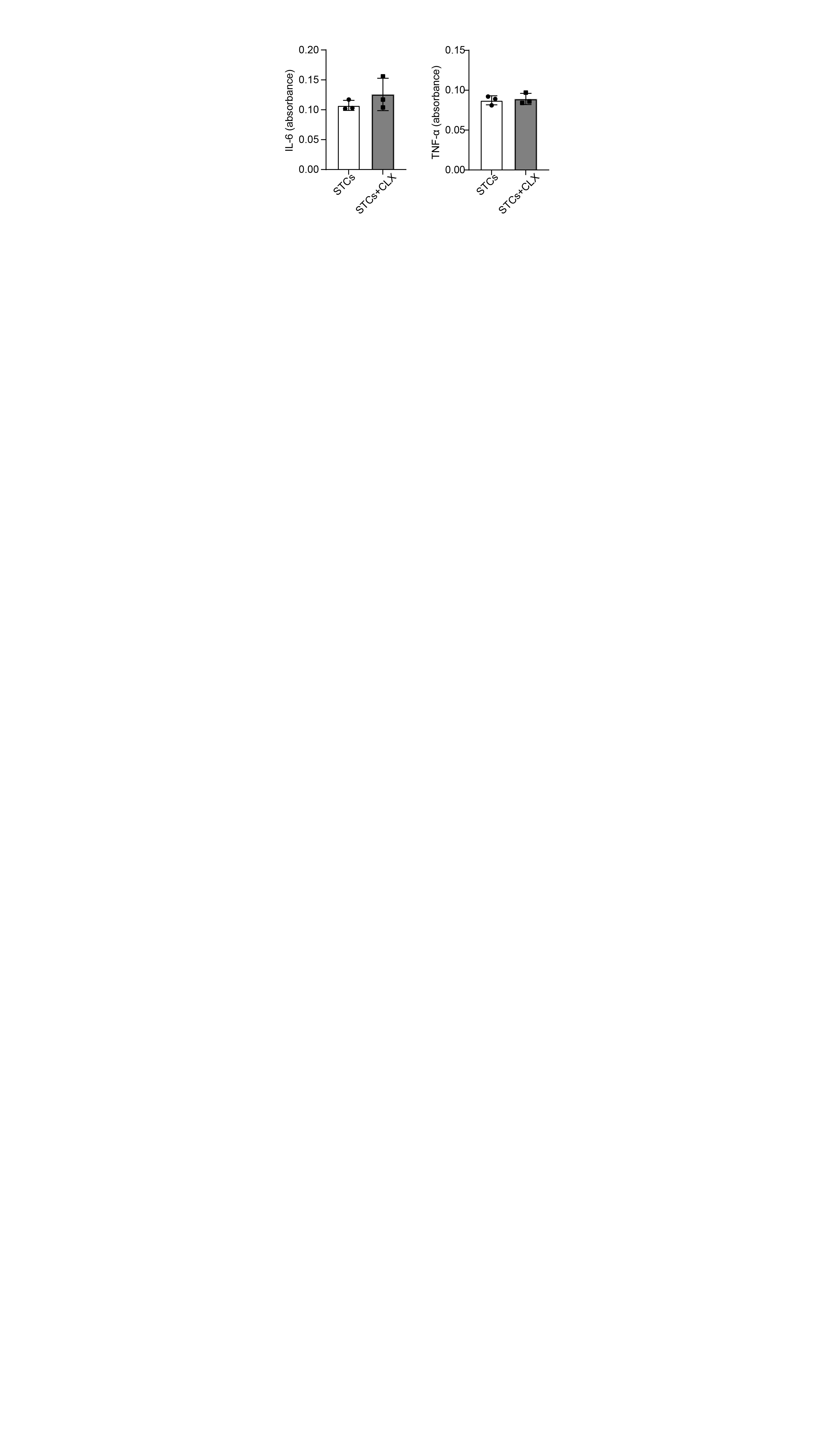

**Figure S5.** IL-6 and TNF-α secretion in senescent B16-F10 cells with or without celecoxib treatment.

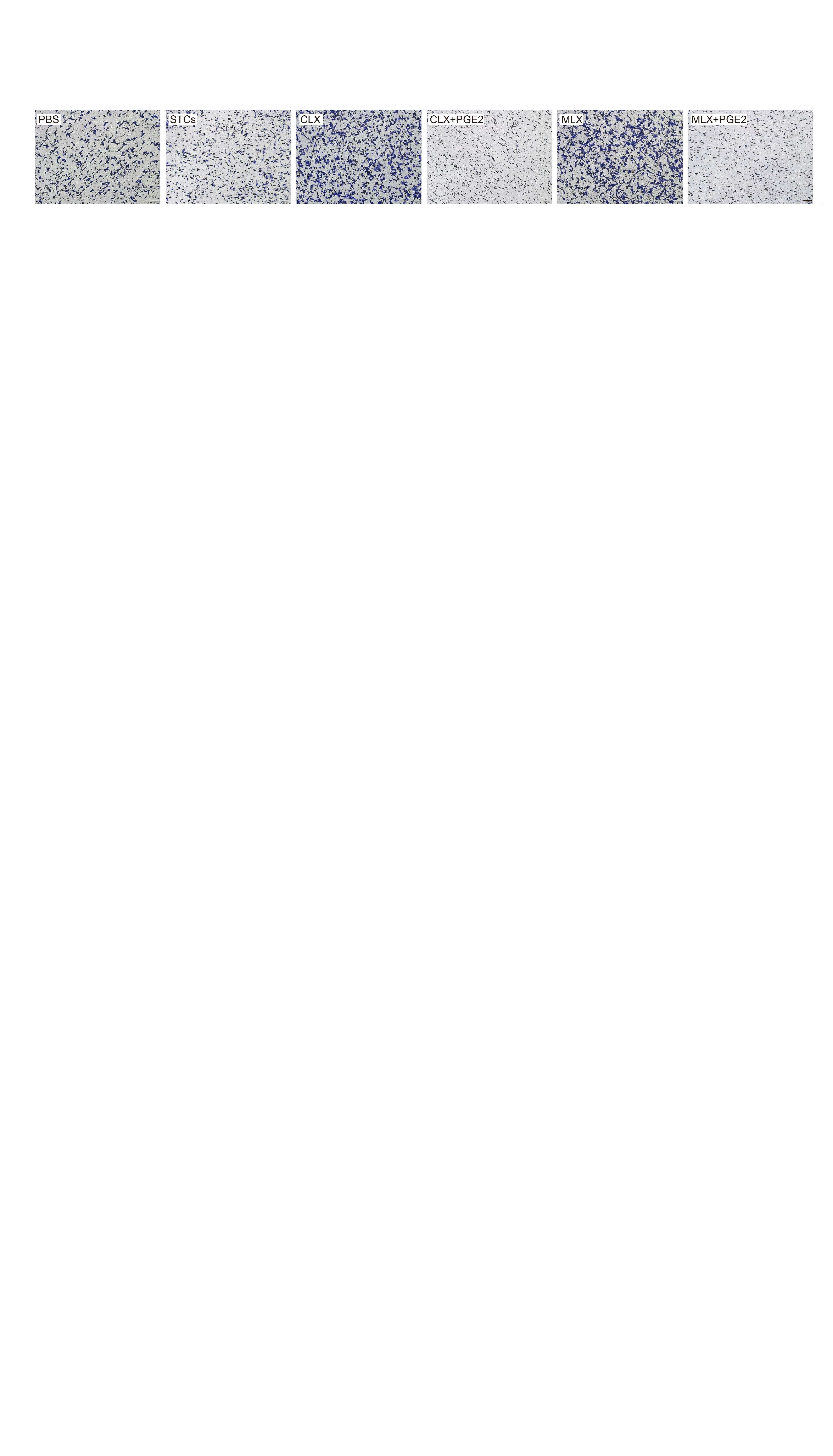

**Figure S6.** Representative images of migrated DCs after different treatments (celecoxib (CLX), meloxicam (MLX), CLX + PGE2, MLX + PGE2, and a vehicle control (PBS)). Both COX2 inhibitors (CLX and MLX) enhanced NK-mediated DC migration compared to PBS or untreated STC groups. Addition of exogenous PGE2 reversed the enhanced DC-recruitment effect of both inhibitors. Scale bar: 200 μm.

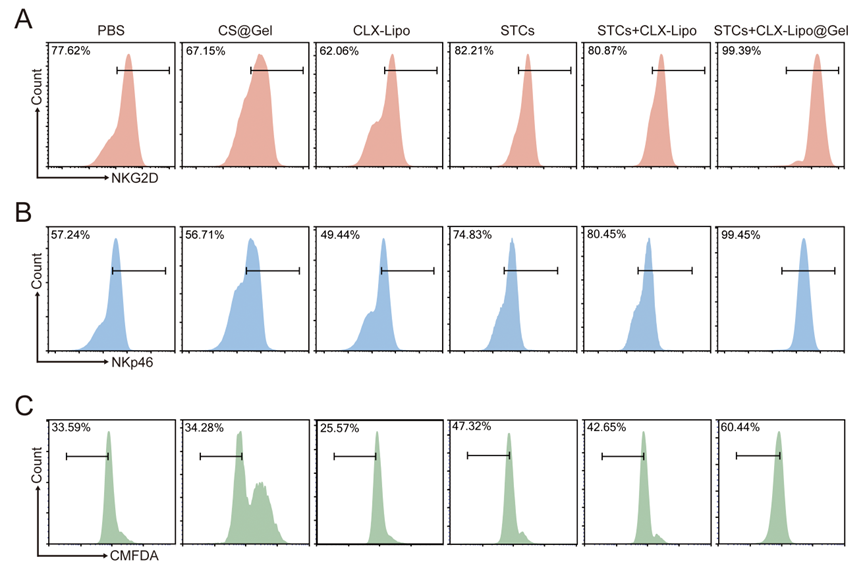

**Figure S7.** Flow cytometry histograms illustrating fluorescence intensity of (A) NKG2D^+^, (B) NKp46^+^, and (C) proliferative NK cells.

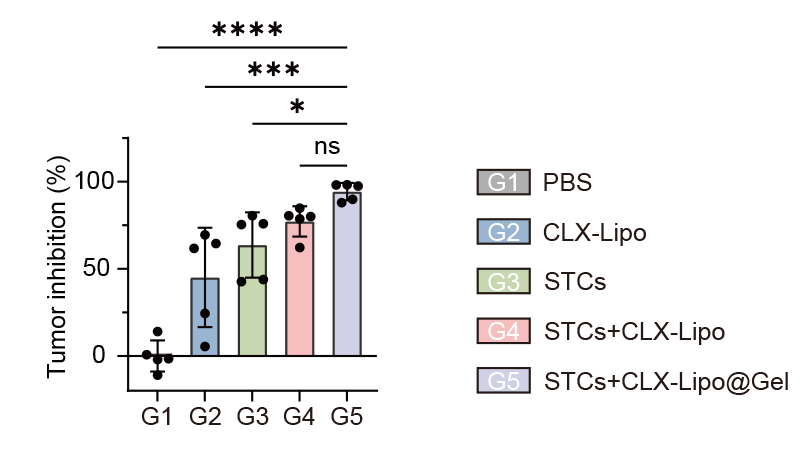

**Figure S8.** Tumor inhibition rate of different groups. Statistical significance was assessed by one-way ANOVA and Tukey multiple comparisons tests. Data were presented as mean ± SD. Statistical significance was determined as ns (not significant, p > 0.05), *p < 0.05, **p < 0.01, ***p < 0.001, ****p < 0.0001.

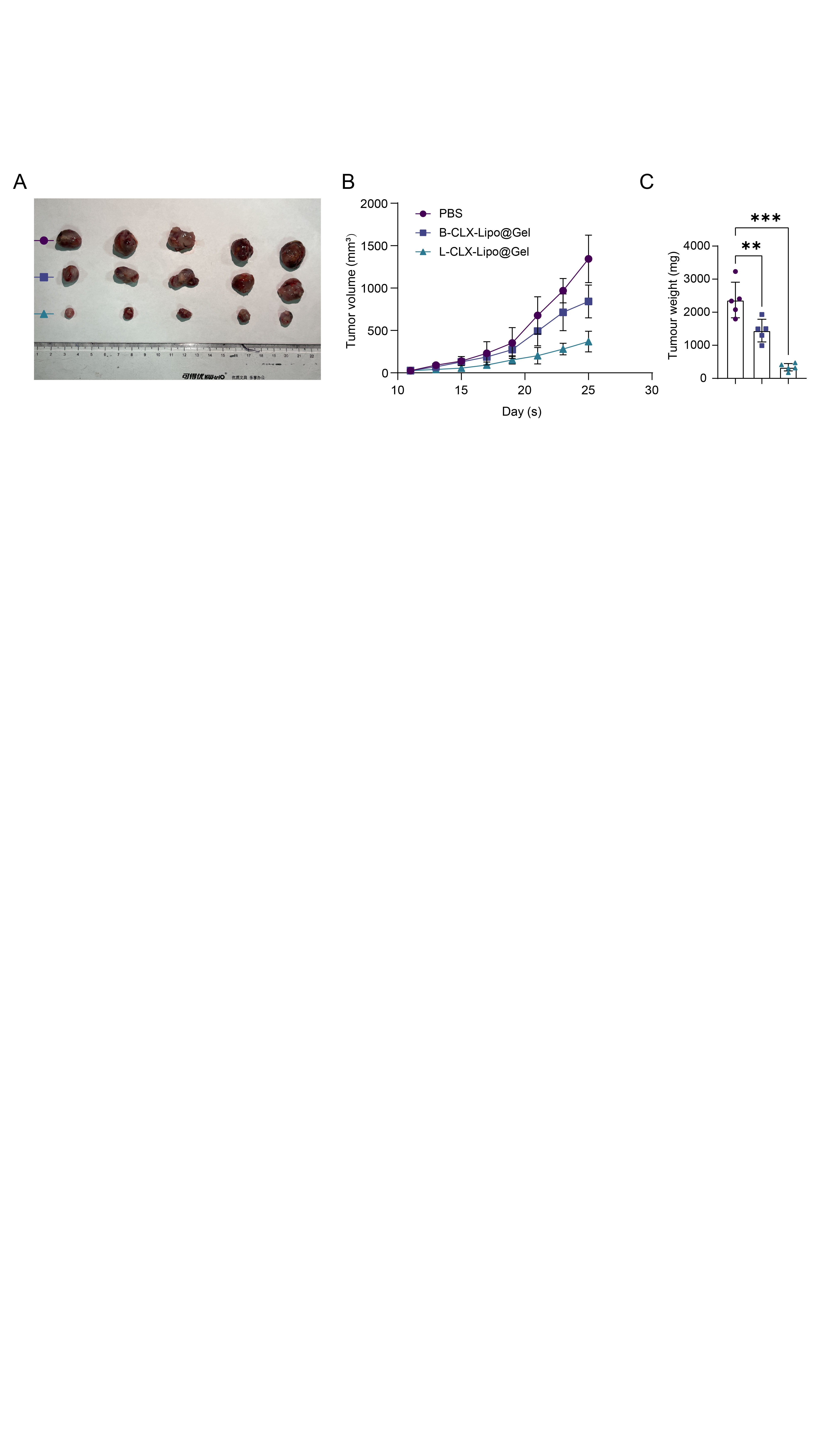

**Figure S9.** Antigen specificity and therapeutic efficacy of the hydrogel vaccine against subcutaneous LLC tumors. (A) Representative photographs of excised tumors. (B) Tumor growth curves of mice during treatment. (C) Weights of excised tumors measured at the endpoint. Data are presented as mean ± SD (n = 5). Statistical significance was analyzed using one-way ANOVA followed by Tukey's post hoc test (ns, *p < 0.05, **p < 0.01, ***p < 0.001). LLC-CLX-Lipo@Gel: senescent LLC cell vaccine; B16F10-CLX-Lipo@Gel: senescent B16F10 cell vaccine.

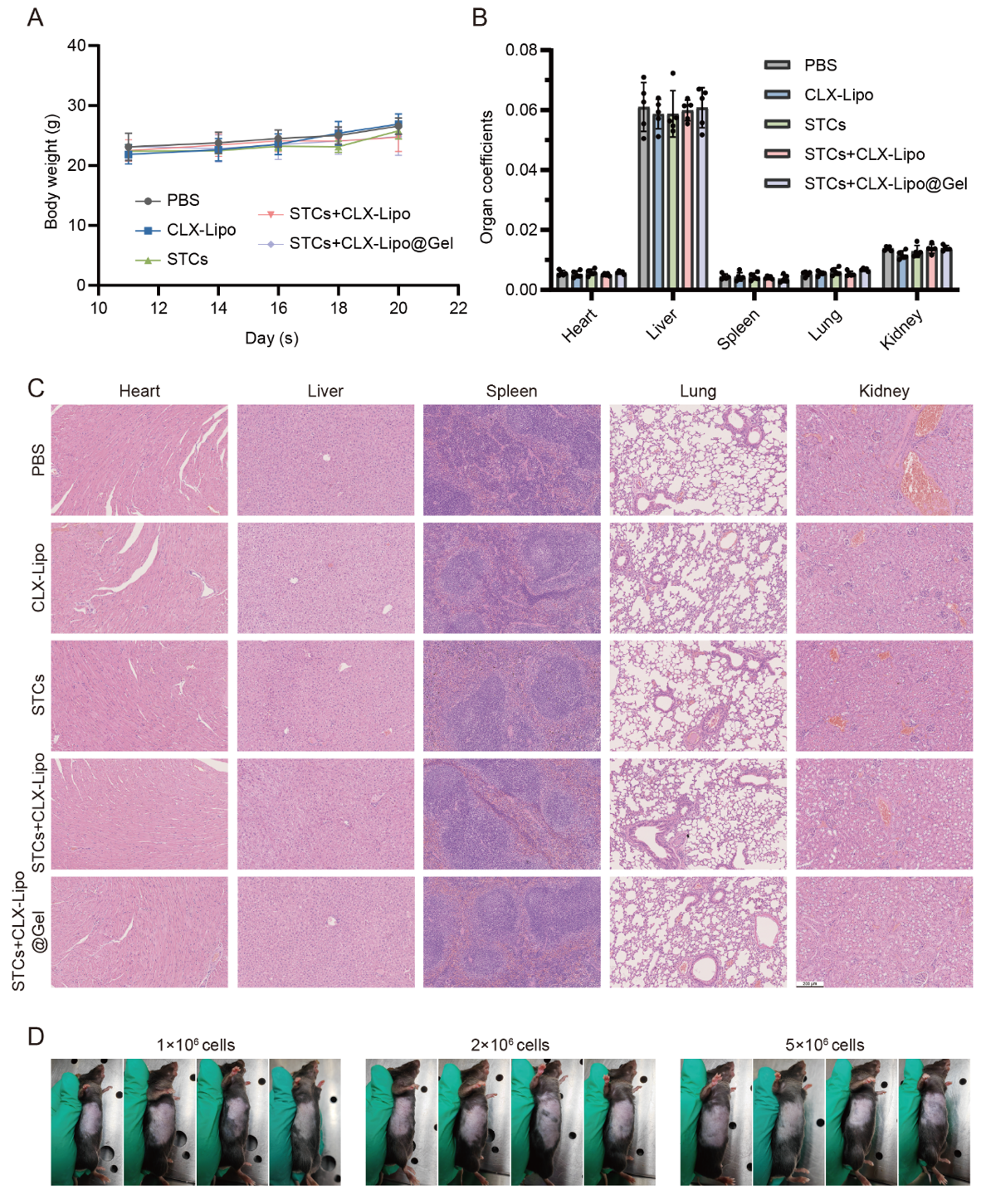

**Figure S10.** Biosafety evaluation of STCs+CLX-Lipo@Gel. (A) Body weight changes during different treatments; (B) Organ coefficients after different treatments; (C) Histological examination of the major organs in each group; (D) Tumorigenic potential of senescent B16-F10 cells. Imaging was conducted on the 42nd day post-subcutaneous administration of varying quantities of senescent B16-F10 cells. Data were presented as mean ± SD (n=5).

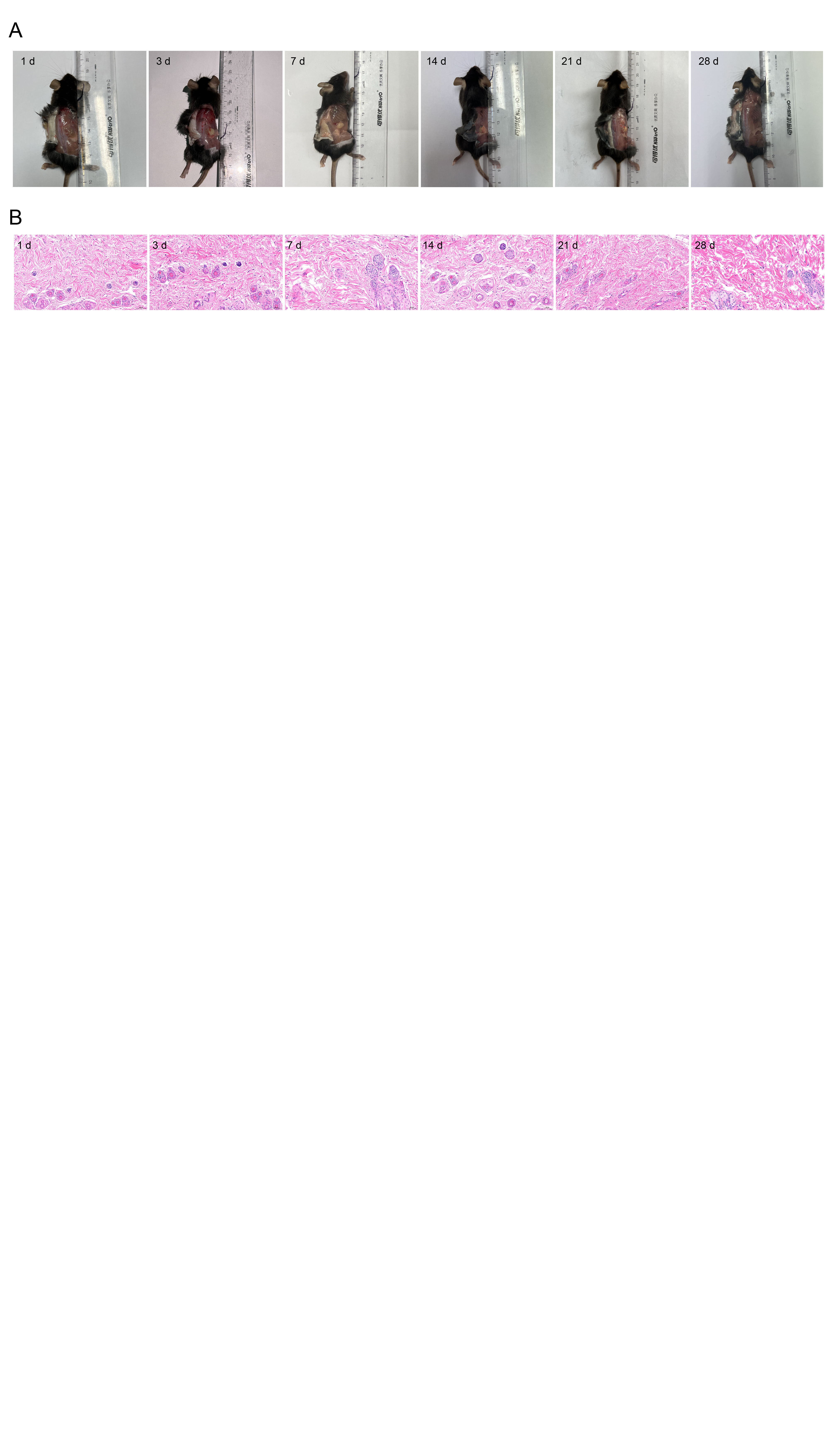

**Figure S11.** Biosafety and degradation evaluation of hydrogel. (A) Hydrogel degradation at different time points; hydrogel completely degraded in day 28. (B) Histological examination of the skin around the injection region. No abnormal immune-cell infiltration or tissue damage.

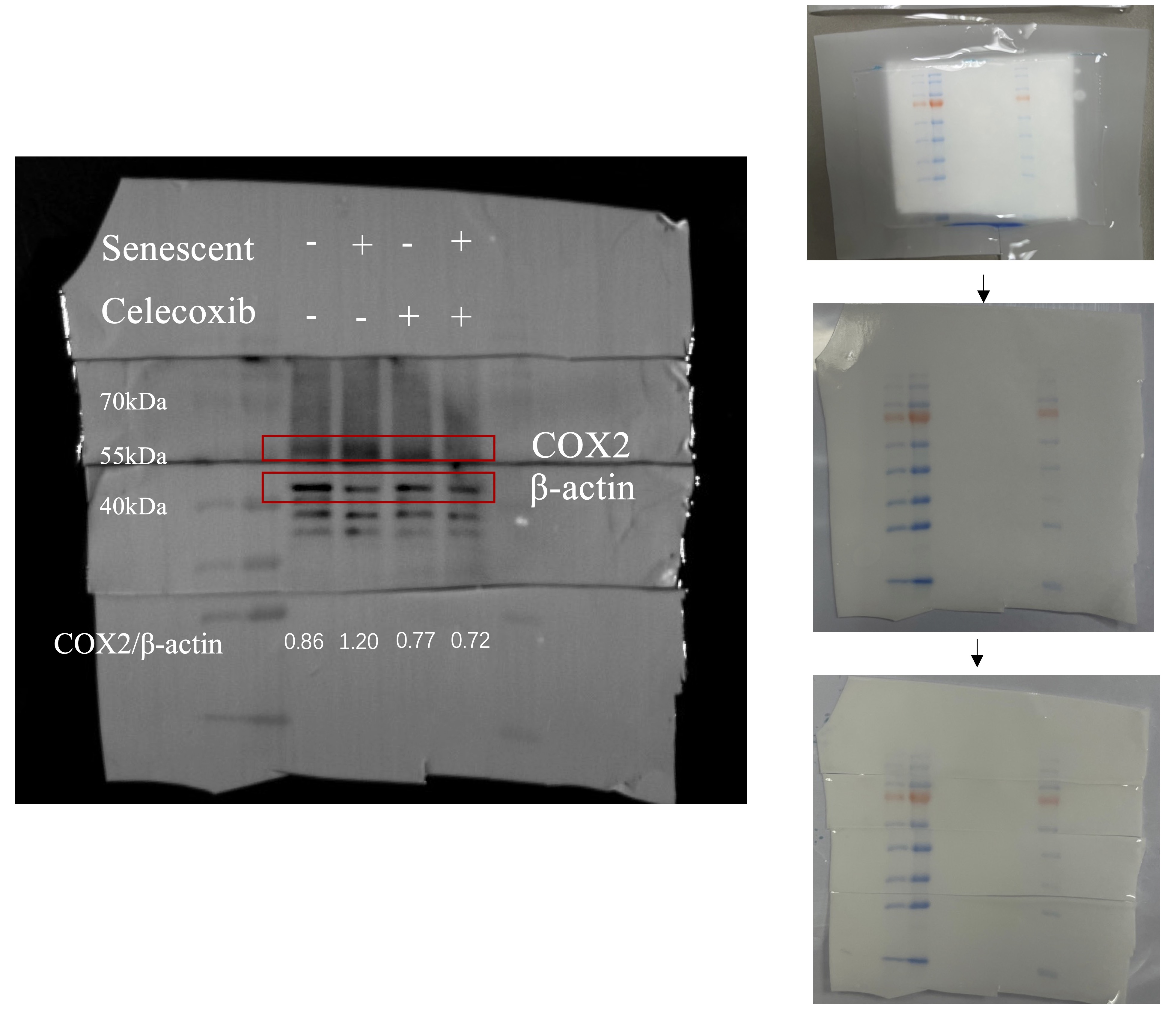

**Figure S12.** Raw data of western blots in Figure 3A.

**Table S1.** Drug encapsulation efficiency and drug-loading capacity of CLX-Lipo.

| Encapsulation efficiency | Loading capacity (%) |
| --- | --- |
| 73.49 ± 1.66 | 2.00 ± 0.04 |

**Table S2.** The gelation time of CS@Gel at different concentrations of gelating agent.

| NaHCO_3_ (M) | PB (M) | Gelation time (min) |
| --- | --- | --- |
| 0.05 | 0.04 | 10 |
| 0.075 |  | 12 |
| 0.1 |  | 45 |

**Table S3.** Primer sequences for qPCR analysis.

| Gene | Forward Primer (5′ to 3′) | Reverse Primer (5′ to 3′) |
| --- | --- | --- |
| *Cdkn1α* | ATCACCAGGATTGGACATGG | CGGTGTCAGAGTCTAGGGGA |
| *CCL2* | ACTGAAGCCAGCTCTCTCTTCCTC | TTCCTTCTTGGGGTCAGCACAGAC |
| *CXCL2* | TCCAGAGCTTGAGTGTGACG | TCAGTTAGCCTTGCCTTTGTTC |
| *IL-1α* | CGCTTGAGTCGGCAAAGAAAT | TGGCAGAACTGTAGTCTTCGT |
| *MMP9* | TCAGGGAGATGCCCATTTCG | GAACGGGAACACACAGGGTTTG |
| *IL-2* | TCCTCACAGTGACCTCAAGTCC | TGACAGAAGGCTATCCATCTCC |
| *IL-12a* | ACGCACTTGTCCTTGAGATG | CTGACCTTGGGAGACACATTT |
| *IL-12b* | TGGGACCTTGGGGTAATGCT | CAGGTGTGGTTGGGCTTCAC |
| *ULBP1* | GTTTGGTAAAGAGGGGAGGGG | AGAAAGGGGCACTAGAAGTTGC |
| *IL-15* | TGCAGCGCGTAAGGAGATG | CGCTCCTTGTGCCTTGATTG |
| *IFN-γ* | AATCAAGCTGCCTCCCGTAT | CCTGCTTCACCTCCTAGCTTT |
| *RAE1* | CTCTGGAGCCAAAAGTCTGAGT | GACTGGCAAGCAGTGGTGTA |
| *CD155* | ACGCAGGCATAAGACGACAA | ATGAGGTAGGAGGGTTGCCT |
| *CCL5* | TGCTGGGCTCTTTACACGAT | TTATGGAGCTTCGGCTTGGG |
| *β-Actin* | CGGTTCCGATGCCCTGAGGCTCTT | CGTCACACTTCATGATGGAATTGA |
